# Diverse branching forms regulated by a core auxin transport mechanism in plants

**DOI:** 10.1101/2022.09.15.508075

**Authors:** Victoria Spencer, Lucy Bentall, C. Jill Harrison

## Abstract

Diverse branching forms have evolved multiple times across the tree of life to facilitate resource acquisition and exchange with the environment. In land plants, sporophyte branching enabled the diversification of the dominant vascular plant clade and arose in their last shared common ancestor; the bryophyte sisters to vascular plants are unbranched. Mechanisms for sporophyte branching are well known in Arabidopsis, where branch initiation and plastic branch outgrowth require directional auxin transport by PIN proteins. However, no broadly applicable genetic mechanisms for branching in vascular plants are known. We have used a combination of surgical and pharmacological treatments and *PIN* expression analyses in the lycophyte *Selaginella kraussiana* to identify PIN-mediated auxin transport as the ancestral mechanism for branching within vascular plants. We show that shortrange auxin transport out of the shoot tips promotes branching, and that branch dominance is coordinated by long-range auxin transport throughout the shoot system. Moreover, the plastic outgrowth of a branch from a unique organ system innovated in lycophytes (the rhizophore) is regulated by long-range auxin transport and associated with a transitory drop in *PIN* expression. We conclude that an ancestral mechanism for branching was independently recruited into plastic branch outgrowth in lycophytes and seed plants. Considered in conjunction with data from other species, our results highlight a pivotal role for the co-option of PINs into the evolution of branching in diverse plant forms.

## Introduction

Branching architectures have evolved many times in different phyla to enable organisms to optimise resource acquisition and exchange with the environment (Harrison, 2017a; Coudert et al., 2019). In land plants, branching forms originated and diversified independently in the haploid gametophyte and diploid sporophyte stages of the life cycle (Harrison, 2017). Whilst gametophyte branching evolved multiple times in the filamentous, thallose or shoot-like forms of bryophytes and ferns, sporophyte branching is thought to have had a single origin in the last common ancestor of vascular plants (Harrison, 2017). In the bryophyte sister clade to vascular plants, sporophytes are unbranched with a single stem terminating in a reproductive sporangium, and eophyte (early land plant) fossils such as *Partitatheca* otherwise resemble bryophyte sporophytes but are branched (Harrison, 2017; Edwards et al., 2021, 2022). The innovation of branching in vascular plants enabled the radiation of diverse shoot architectures and led to a tenfold increase in plant species numbers (Harrison and Morris, 2018). Thus, identifying mechanisms enabling the origin and diversification of vascular plant branching patterns is a key goal of evolutionary biologists.

Most of our knowledge of the genetic mechanisms regulating sporophyte branching was generated in flowering plants such as Arabidopsis, where short-range PIN-mediated auxin transport away from the leaf axils generates auxin minima and enables the establishment of axillary meristems (Q. Wang et al., 2014; Y. Wang et al., 2014). Axillary meristems can remain dormant for long periods, and their outgrowth as branches in different parts of the shoot system is globally coordinated by long-range PIN-mediated auxin transport in the stems (Müller and Leyser, 2011). Active shoot apices act as auxin sources, exporting auxin basipetally via the polar auxin transport stream of the stem vasculature (Thimann and Skoog, 1933). This basipetal auxin flow blocks the capacity for auxin export from dormant axillary meristems (Prusinkiewicz et al., 2009). However, if basipetal auxin transport is disrupted, e.g. by excision of the shoot apices, the axillary buds can export auxin, enabling branch outgrowth (Thimann and Skoog, 1933; Prusinkiewicz et al., 2009).

Whilst these roles for PIN-mediated polar auxin transport in branch initiation and outgrowth are well known in Arabidopsis, mechanisms for branching are poorly understood in other plant groups (Harrison, 2017a; Coudert et al., 2019). In the gametophytic leafy shoots of a moss (*Physcomitrium patens*), recent work identified roles for diffusive auxin transport from an apical auxin source in the regulation of apical dominance and branching, but moss leafy shoot branching is analogous to axillary branching in seed plants and has a different cellular basis (Coudert et al., 2015; Thelander et al., 2022). The filamentous tissues of *P. patens* are also branched, with dominance exerted by the apical cell of each filament (Viaene et al., 2014; Coudert et al., 2019; Nemec-Venza et al., 2022). Here, PIN-mediated auxin transport out of the apical cells suppresses a foraging growth habit with strong apical dominance, giving plants a more uniform, circular form (Viaene et al., 2014; Nemec-Venza et al., 2022). In *P. patens* sporophytes, disruption of PIN-mediated polar auxin transport can induce branching, leading plants to resemble the earliest sporophytic branching forms in the fossil record (Bennett, Liu, et al., 2014; Edwards et al., 2014; Harrison and Morris, 2018). Whilst these data implicate auxin transport in the evolution of diverse branching forms, they were generated using a single species, and mosses are distantly related to Arabidopsis and other seed plants (Puttick et al., 2018). Moreover, PIN proteins diversified independently in bryophytes and vascular plants; vascular plant PINs are thought to have originated from a single canonical ancestral PIN and to have diversified independently in lycophytes and euphyllophytes (Bennett, Brockington, et al., 2014).

Lycophytes are a key group to resolve questions about the evolution of sporophyte branching in land plants (Spencer et al., 2021). They originated over 420 million years ago (Morris et al., 2018), and some species closely resemble their fossil relatives, exhibiting nascent branching architectures (Harrison and Morris, 2018). Lycophytes are the sister group to euphyllophytes (ferns and seed plants) and are hence ideally placed in the plant tree of life to identify vascular plant homologies (Spencer et al., 2021). Lycophytes branch by bifurcation (Gola, 2014), which is the ancestral pattern of branching within vascular plants. The cellular basis of bifurcation in the lycophyte *Selaginella kraussiana* was resolved through clonal analysis and shown to involve cyclical expansion of the apical cell pool, broadening of the parent shoot apex, and the segregation of apical cells to form two new shoot apices before branches diverge (Harrison et al., 2007; Harrison and Langdale, 2010). Whereas major branches initiate from two apical cells, minor branches initiate from a single apical cell (Harrison et al., 2007; Harrison and Langdale, 2010). A unique leafless organ system of Selaginellales (the rhizophore) can also contribute to the overall pattern of branching (Banks, 2009; Spencer et al., 2021). Rhizophores develop from angle meristems initiated at sites of branch divergence, have gravitropic growth, and normally start to develop roots when they reach the soil. However, if the shoot tips above an angle meristem are excised, the rhizophore can instead develop a branch (Banks, 2009; Spencer et al., 2021). Prior surgical and pharmacological experiments in *Selaginella* identified roles for polar auxin transport in the outgrowth of branches from angle meristems, but a range of species and experimental designs were used (Williams, 1937b; Wochok and Sussex, 1975; Jernstedt et al., 1994; Mello et al., 2019), so it is not clear how broadly applicable inferences are. Auxin has been shown to localize in the stem vasculature of *S. wildenowii* (Wochok and Sussex, 1973), there is long range basipetal transport in *S. kraussiana* stems (Sanders and Langdale, 2013), and auxin transport inhibition leads to shoot apex termination in *S. kraussiana* (Sanders and Langdale, 2013). As plastic branching responses evolved independently in lycophytes and seed plants, we chose *S. kraussiana* as a model system to explore mechanisms of bifurcation in the origin of vascular plants and the independent evolution of plasticity.

Here we report that auxin transport in *S. kraussiana* promotes and coordinates bifurcation throughout the shoot system, and that basipetal auxin transport from the shoot apices suppresses branch outgrowth from angle meristems. Of four *S. kraussiana PINs* (*SkPINs*), two (*SkPINR* and *SkPINS*) are expressed in the shoot tips, and three (*SkPINR, SkPINS* and *SkPINT*) in the stem vasculature. Bifurcation is sensitive to pharmacological inhibition of PIN function, and branch outgrowth from angle meristems follows a drop in *SkPINR* and *SkPINS* expression. We conclude that PIN-mediated auxin transport is an ancestral regulator of vascular plant branching that was independently co-opted into the evolution of plastic branching responses in lycophytes and seed plants.

## Results

### Bifurcation and rhizophore emergence proceed consistently during development

*S. kraussiana* has sprawling prostrate shoots (Fig. 1A), which bifurcate through unequal dichotomy to produce larger “major” branch and smaller “minor” branches (Fig. 1B; arrows, Fig. 1C-D). Due to the left-right alternation of major and minor branches, a zig-zig like architecture is produced (Fig. 1E, I). At each bifurcation point (BP) (arrowhead, Fig.1D, F) Angle Meristems (AMs) subsequently initiate, and these typically produce a rhizophore (Fig. 1G). However, angle meristems have plastic identity and in some instances can produce branches (Fig. 1H). To provide a baseline for comparison of branching phenotypes following experimental interventions, we first characterised patterns of apex bifurcation and angle meristem outgrowth in explants grown on soil (Fig. 1). Shoot explants with four bifurcation points were propagated as cuttings, and the number of bifurcation points along the main axis was recorded in an 8-week time course (Fig. 1J). The developmental time interval (plastochron) between successive bifurcations along the main axis ranged from < 1 to > 4 weeks and the mean was 1.6 weeks (Supplementary Fig. 1). The timing of rhizophore emergence from each angle meristem was determined by calculating the percentage of bifurcation points bearing a rhizophore each week (Fig. 1E, K). At week 0, the most basal angle meristems (BP1) had all produced a rhizophore, but this percentage decreased with proximity to the leading apex (Fig. 1E, K). During subsequent weeks, the angle meristems produced rhizophores in turn such that by week 8, the five most basal bifurcation points all bore rhizophores (BP1-5, Fig. 1K). Thus, bifurcation and ensuing rhizophore emergence proceed in predictable patterns in *S. kraussiana*.

**Figure 1:**
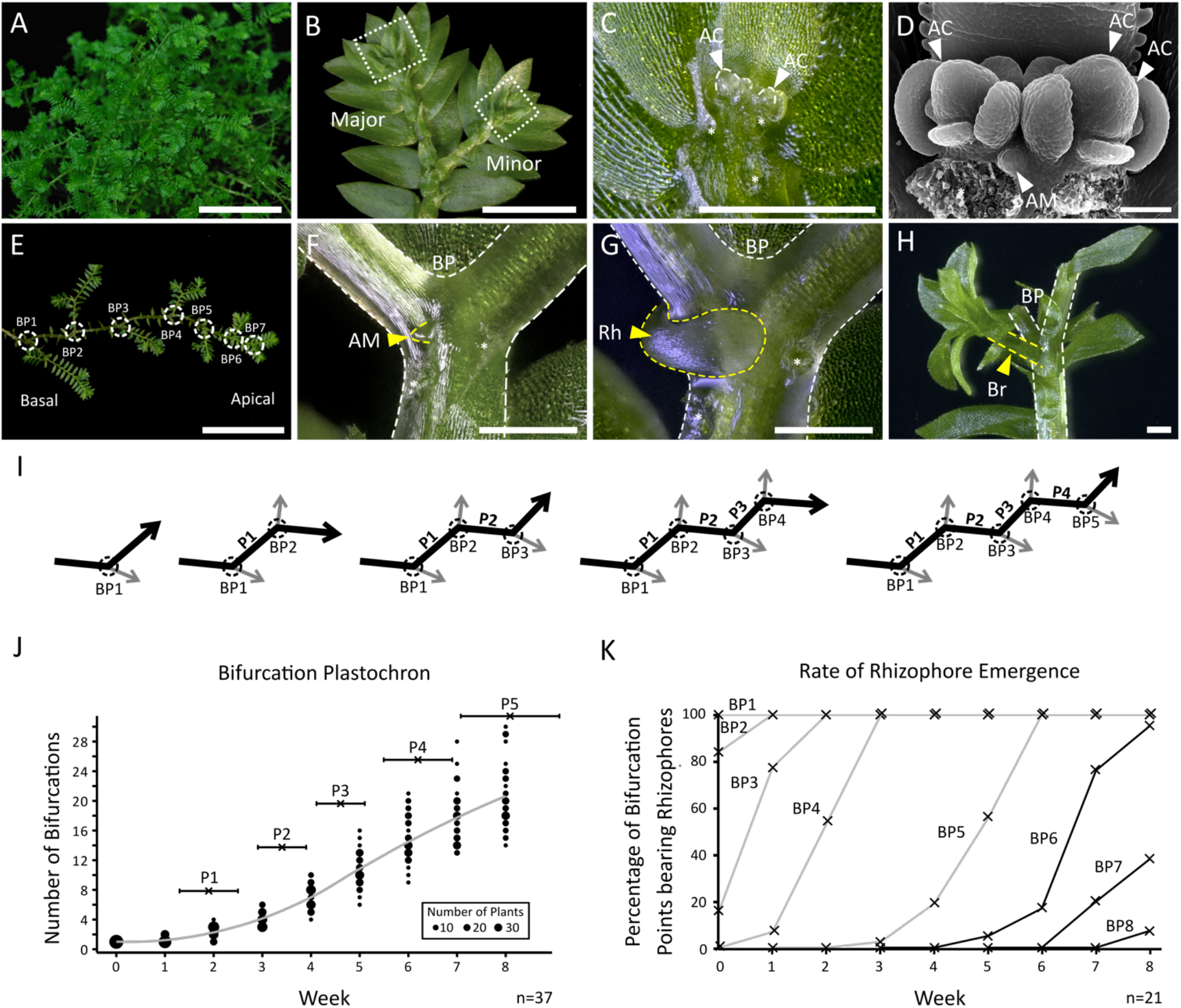
Branching properties of the lycophyte, *Selaginella kraussiana*. (**A**) Overall growth habit of *S. kraussiana*, showing its creeping prostrate shoot system with multiple shoot apices. Scale bar = 5 cm. (**B**) Light micrograph of *S. kraussiana* apices, showing anisotomous bifurcation to produce major and minor branches. The white dotted boxes in **B** indicate shoot apices dissected and magnified in images **C** and **D**. Scale bar = 5 mm. (**C**) Light micrograph of *S. kraussiana* apices with dorsal leaves removed to reveal the meristems. Asterisks show the position of dissected leaf scars. AC = apical cells. Scale bar = 0.5 mm. (**D**) Scanning Electron Micrograph of *S. kraussiana* apices. AC = apical cells; AM = angle meristem. Asterisks show the position of dissected leaf scars. Scale bar = 50 μm. (**E**) Image of a *S. kraussiana* shoot illustrating successive bifurcation point (BPs; white circles) and strong apical dominance. Scale bar = 5 cm. (**F**-**H**) Light micrograph of (**F**) a dorsal angle meristem, (**G**) an emerging rhizophore, and (**H**) an emerging branch at a bifurcation point (BP). AM = angle meristem; Rh = rhizophore; Br = branch. Scale bars = 0.5 mm. (**I**) Schematic illustrating the pattern of bifurcation in *S. kraussiana* shoots, with successive events generating major (black) and minor (grey) branches. The time interval between bifurcations was used to define the bifurcation plastochron (P1-P4), and arrows represent actively growing apices. (**J**) Graph representing the bifurcation plastochron (P1-P5) of *S. kraussiana* cuttings grown on soil in an eight-week time course. Dot size represents the frequency of plants with a corresponding number of total bifurcations on the y axis, whilst the fitted line shows a local regression. N = 37. (**K**) Percentage of bifurcation points bearing rhizophores in an eight-week time course. At week 0, bifurcation points at the base of each explant (BP1) bore a rhizophore. By week 8, each plant bore rhizophores at the five most basal bifurcation points (BP1-5). N =21.

### Auxin suppresses and auxin transport promotes bifurcation

To identify potential roles for auxin and PIN-mediated auxin transport in *S. kraussiana* branching, we first evaluated the effects of exogenously applied auxin (1-Napthaleacetic acid; NAA) and auxin transport inhibitors (N-1-Naphthylphthalamic Acid; NPA) on shoot explants grown for eight weeks in axenic culture (Fig. 2, Supplementary Fig. 2). This showed a general suppressive effect of NAA on growth as evaluated by the length of the main axis (Fig. 2A, B). There were also dose dependent decreases in the number of main axis plastochrons and the overall number of bifurcations, but no effects of NAA on the length of individual plastochrons (data for plastochron 1 shown) or on the number of leaves per plastochron were discernible (Fig. 2B). Thus, shoot development proceeds at a slower rate than normal following NAA treatment. From these data we concluded that NAA has a suppressive effect on bifurcation and, as bifurcation reflects the activity of apical cells in the shoot apex (Harrison et al., 2007; Harrison and Langdale, 2010; Sanders and Langdale, 2013), that auxin regulates apical cell activity. As explants grown with NPA showed a similar response to explants grown with NAA (Fig. 2C, D), and auxin transport out of apical cells is required for their function (Sanders and Langdale, 2013), we infer that auxin accumulates in the apical cells of NPA treated plants. This accumulation suppresses bifurcation, hence, short-range auxin transport out of the apical cells is likely to promote bifurcation.

**Figure 2:**
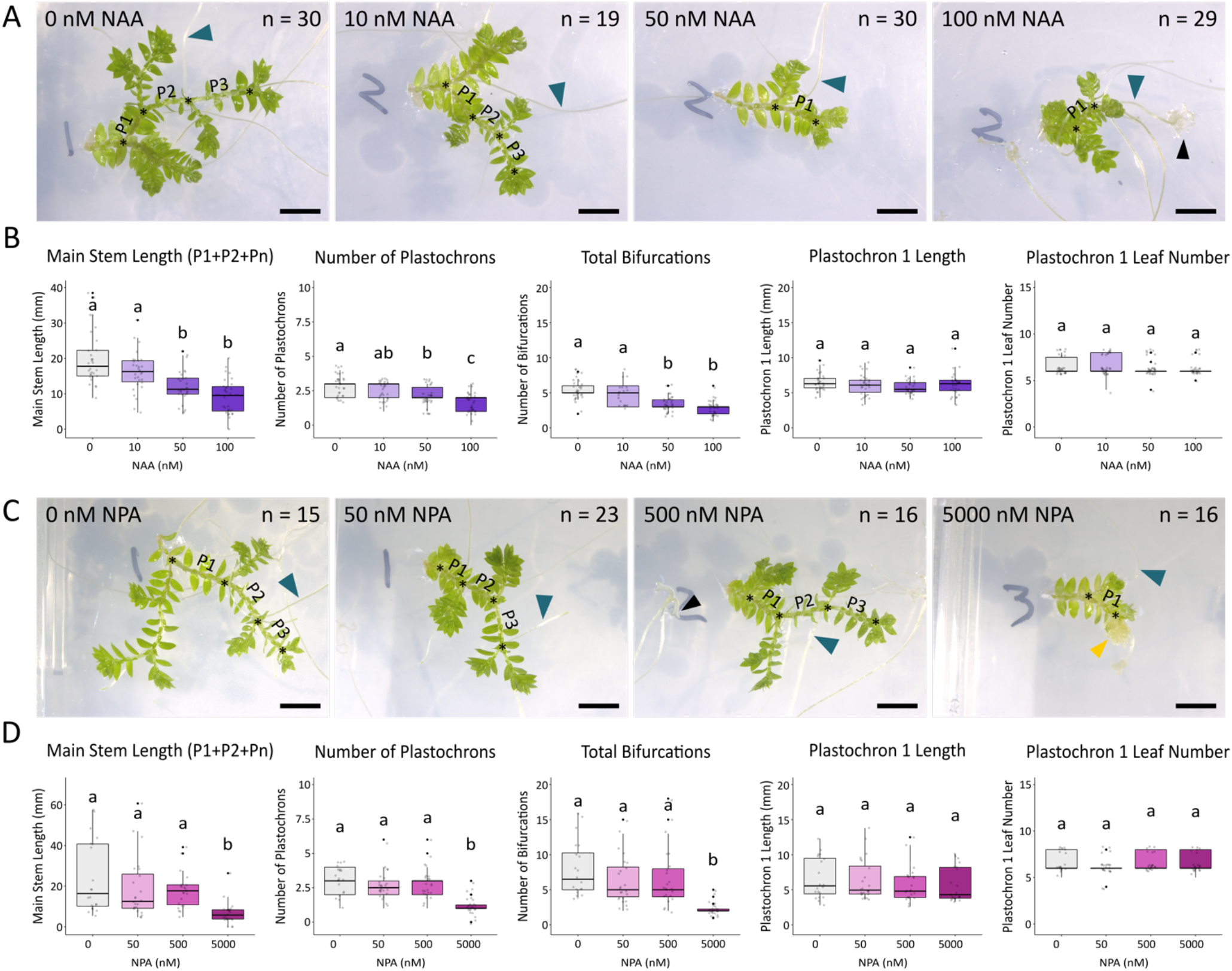
Auxin and short-range auxin transport regulate bifurcation. (**A**) Images of *S. kraussiana* explants grown for eight weeks in tissue culture media supplemented with 0 nM, 10 nM, 50 nM and 100 nM NAA. Black asterisks mark bifurcation points and P numbers indicate plastochrons of the main axis. Blue arrowhead = rhizophore, black arrowhead = root. Scale bar = 5 mm. (**B**) Graphs showing that NAA treatment suppresses bifurcation without affecting plastochron length or leaf initiation. N = 19-30. In all graphs, boxes represent lower quartile, median and upper quartile. Lines represent spread of data and black points show outliers. One-way ANOVA tests with Tukey or Kruskal Wallis multiple comparisons were performed (F(3,116)= 20.44, p<0001; F(3,116)= 18.9, p<0.0001; F(3, 104)= 26.21, p<0.0001; F(3,115)= 1.33, p=0.27; χ^2^(3)= 2.01, p=0.57). (**C**) Images of *S. kraussiana* explants grown for eight weeks in tissue culture media supplemented with 0 nM, 50 nM, 500 nM and 5 μM NPA. Black asterisks mark bifurcation points and P numbers indicate plastochrons of the main axis. Blue arrowhead = rhizophore, black arrowhead = root, yellow arrowhead = callus. (**D**) Graphs showing that NPA treatment suppresses bifurcation without affecting plastochron length or leaf initiation. N = 15-23. In all graphs, boxes represent lower quartile, median and upper quartile. Lines represent spread of data and black points show outliers. One-way ANOVA tests with Tukey or Kruskal Wallis multiple comparisons were performed (F(3,87)= 7.07, p=0.00026; F(3,87)= 16.6, p<0.0001; F(3,87)= 12.22, p<0.0001; F(3,84)= 0.59, p=0.62; χ^2^(3)= 3.05, p=0.38). Data from one of three experimental replicates are shown.

### Long range auxin transport mediates apical dominance and co-ordinates overall bifurcation patterns

To identify any long-range effects of auxin and auxin transport on bifurcation, we designed and implemented a series of surgical and pharmacological experiments on explants grown on soil (Fig. 3, Supplementary Fig. 3). These explants had three bifurcation points, a leading major branch and three minor branches, respectively Ma and Mi1-3 in Fig. 3A. To identify any long-range dominance effects, the major branch and/or the leading minor branch (minor branch 1; Mi1), the second side branch (minor branch 2; Mi2) or the third side branch (minor branch 3; Mi3) was removed (Fig. 3A), and the number of bifurcations of remaining branches was quantified following eight weeks of growth (Fig. 3B). Whilst there was no effect of minor branch excision on major branch bifurcation, excision of the major branch tips resulted in an increase in bifurcation in minor branches 1 and 2 but not minor branch 3 (Fig 3B). However, when minor branch 1 was excised as well as the major branch, the number of bifurcations of minor branch 3 increased significantly (Fig 3B). These data suggest that bifurcation involves long-range basipetal movement of a suppressive signal from the branch tips, that the major branches are the strongest source of the signal, that minor branches are also a source, and that the signal does not move acropetally. To test the hypothetical identity of this long-range signal as auxin, surgical experiments were combined with pharmacological experiments applying NAA in lanolin paste to excision sites (Fig. 3C). NAA applied at the major branch tip suppressed bifurcation of minor branch 1 and minor branch 2 but had a weaker effect on minor branch 3, consistent with an identity of the basipetal signal as auxin (Fig. 3D).

**Figure 3:**
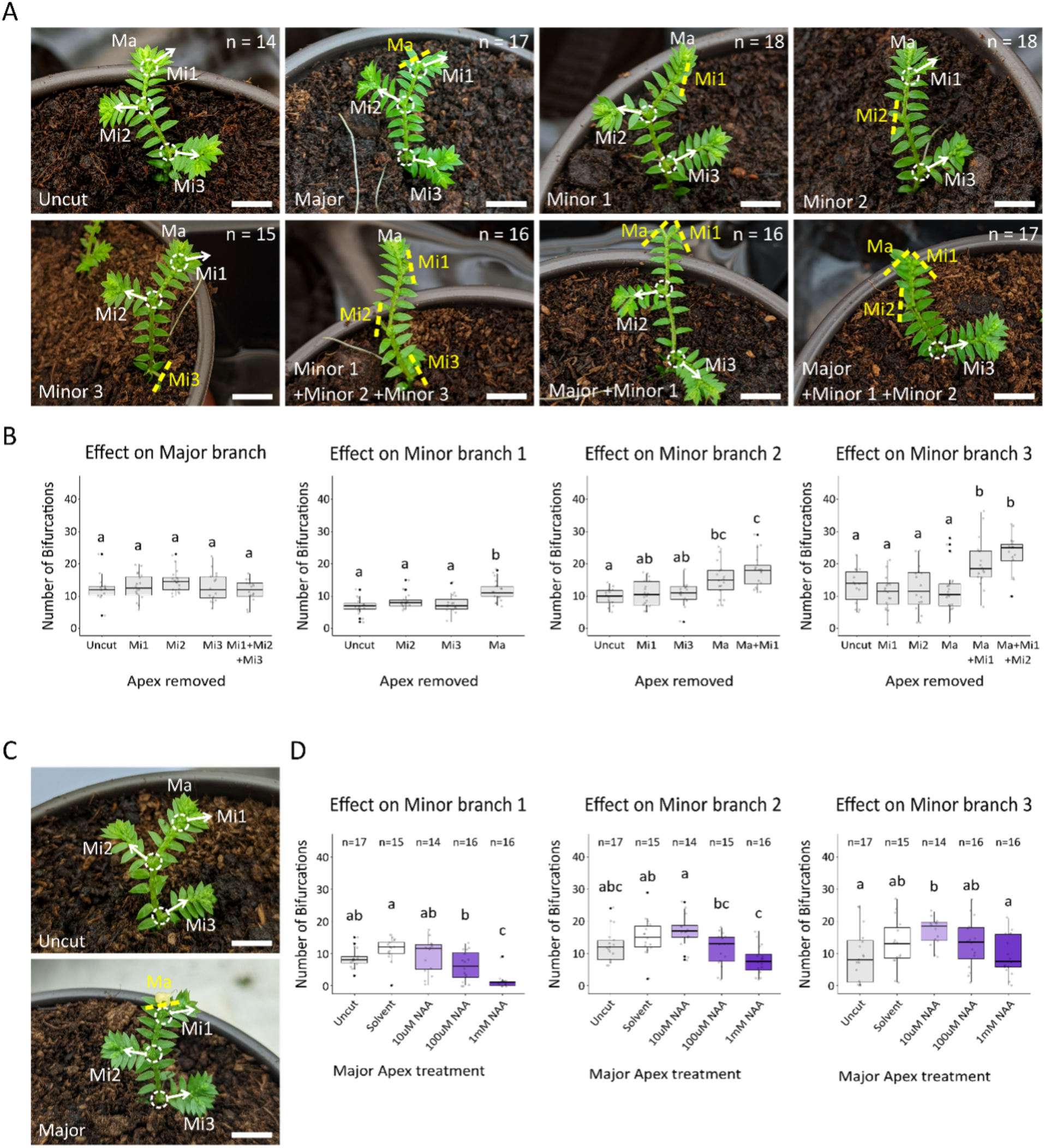
Long-range auxin transport co-ordinates bifurcation. (**A**) Photographs illustrating the design of surgical experiments. Explants were left uncut, or the major (Ma), first minor branch (Mi1), second minor branch (Mi2) or third minor branch (Mi3) was removed singularly or in combination, and plants were grown for 8 weeks. Dashed lines indicate excision sites, and dashed circles with arrows indicate intact minor branches. Scale bar = 5 mm. (**B**) Graphs showing the effect of apex removal on bifurcation of the major branch and minor branches 1, 2 and 3. Whilst minor branch removal had no effect on major branch bifurcations, major branch removal affected the activity of minor branches 1 and 2, and jointly with minor branch 1 affected minor branches 2 and 3. N = 15-18. One-way ANOVAs with Tukey tests for multiple comparisons were performed (F(4,76)= 1.28, p=0.28; F(3,60)= 9.83, p<0.0001; F(4,75)= 9.94, p<0.0001; F(5,91)= 8.95, p<0.0001). (**C**) Images of explants with three bifurcation points showing major branch removal and replacement with lanolin paste. Scale bar = 5 mm. (**D**) Graphs showing the number of bifurcations of minor branch 1, 2 and 3 following excision and replacement of the major branch with lanolin paste containing a solvent control or auxin (10 μM NAA, 100 μM or 1 mM NAA) and eight subsequent weeks of growth. With increasing concentrations, auxin progressively inhibited bifurcations at minor branch 1 and 2. In all graphs, boxes represent lower quartile, median and upper quartile. Lines represent spread of data and black points show outliers. One-way ANOVAs with Tukey tests for multiple comparisons were performed (F(4,73)= 14.63, p<0.0001; F(4,72)= 6.86, p<0.0001; F(4,73)= 2.91, p=0.027). Data from one of three experimental replicates are shown.

### Auxin and auxin transport regulate plasticity and branch emergence from angle meristems

A further component of the overall *Selaginella* branching habit involves plastic branch outgrowth from the angle meristems following surgical decapitation. However, previously reported results have used diverse species and experimental systems to evaluate the role of long-range cues in this process (Williams, 1937; Wochok and Sussex, 1975; Jernstedt et al., 1994; Mello et al., 2019). To identify any roles for long-range and polar auxin transport in branch outgrowth from *S. kraussiana* angle meristems, we performed surgical and pharmacological experiments using tissue culture and whole plant systems (Fig. 4, Supplementary Fig. 4). In the tissue culture system, explants with 1 bifurcation point were grown for 5 weeks and their apices were left intact (uncut) or excised (cut) (Fig. 4 A, B). Whereas all angle meristems developed rhizophores after 2 weeks in ‘uncut’ explants, there was a delay in rhizophore emergence in ‘cut’ explants (Fig. 4B). Furthermore, 20% of angle meristems developed to form branches in ‘cut’ explants, consistent with a role for long-range signaling in branch emergence (Fig. 4B). To identify any roles for auxin and auxin transport in branch outgrowth, explants were grown on media containing a range of concentrations of NAA (0 nM, 10 nM, 50 nM, 100 nM), NPA (0 nM, 50 nM, 500 nM, 5 μM) or NAA and NPA in combination (10nM NAA + 50nM NPA), and angle meristem identity was recorded at week 5 (Fig. 4C). Regardless of the concentration of NAA or NPA, ‘uncut’ explants showed no branch outgrowth, but with 5 μM NPA, explants developed a callus like tissue at low frequency (Fig. 4C). In contrast, ‘cut’ explants showed progressive inhibition of branch emergence and restoration of rhizophore production with increasing concentrations of NAA and NPA (Fig. 4C). Thus, we infer that auxin generated in the shoot tips acts on rhizophore identity normally repressing branch outgrowth, and that decapitation releases this repression.

**Figure 4:**
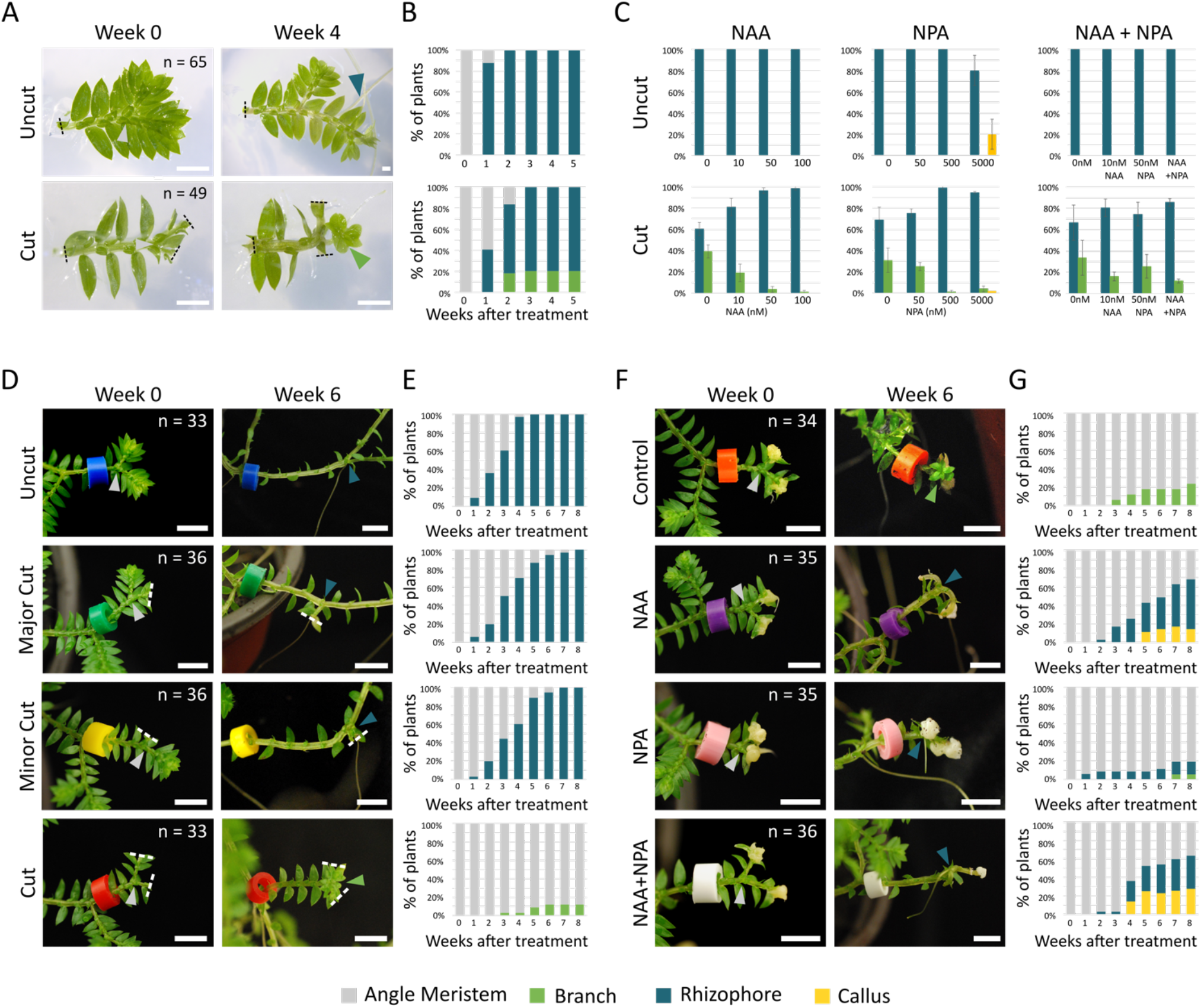
Apical auxin inhibits branch formation from the Angle Meristem (AM). (**A**) Images showing sterile *S. kraussiana* explants with one bifurcation point grown for five weeks in tissue culture. Explants either had their shoot tips left intact (uncut) or excised (cut), and uncut explants developed rhizophores (blue arrowhead) by week two. Cut explants developed rhizophores or, at a low frequency, branches (green arrowhead). Scale bar = 2 mm. (**B**) Graphs showing the frequency of rhizophore and branch development from explants shown in **A**. (**C**) Explants with one bifurcation point were transferred to media containing auxin (0 nM, 10 nM, 50 nM and 100 nM NAA), an auxin transport inhibitor (0 nM, 50 nM, 500 nM, and 5 μM NPA) or a combination (0 nM, 10 nM NAA, 50 nM NPA and 10 nM NAA + 50 nM NPA). The identity of organs produced from the angle meristem was recorded at week 5. NAA and NPA had no effect on angle meristem identity in uncut explants, but in explants with cut apices increasing concentrations led to progressive reductions in branch formation and increased frequencies of rhizophore initiation. High concentrations of NPA sometimes promoted callus formation in cut and uncut explants. Pooled week 5 data from three experimental replicates are shown (individual experiments and timecourses are shown in Supplementary Figure 4). (**D**) Plants with four bifurcation points were grown on soil and were left uncut, had both apices cut, or had either the major or minor branch cut (excisions = dashed white lines). By week 6, rhizophores were produced by all the uncut, major cut and minor cut plants. When both apices were removed, no rhizophores initiated and 12 % of plants produced branches instead. Grey arrows = angle meristems; blue arrows = rhizophores; green arrow = branch. Scale bar = 5 mm. (**E**) Graphs showing the frequency of rhizophore and branch development from explants shown in **D**. (**F**) Images of plants whose apices were removed and replaced with lanolin paste mixed with either a solvent control, 1 mM NAA, 500 μM NPA, or 1 mM NAA + 500 μM NPA. Whilst branches formed in control plants, NAA, NPA and NAA + NPA treatments partially restored rhizophore production or led to production of a callus-like tissue. Auxin treatments also caused stem curling and thickening, and an increase in the divergence angle between branches. Grey arrows = angle meristems; blue arrows = rhizophores; green arrow = branch. Scale bars = 5 mm. (**G**) Graphs showing the frequency of rhizophore and branch development from explants shown in **F**. Data from one of three experimental replicates are shown.

### Long-range cues regulate angle meristem activity, branch outgrowth and rhizophore development

To corroborate findings from the tissue culture system and discern any dominance effects of major and minor branch apices on branch emergence from angle meristems, we next implemented surgical and pharmacological experiments in cuttings with four bifurcation points grown for four weeks on soil (Fig. 4D, E). The major axis of each plant was tagged and either (1) both apices were left intact, ‘uncut’, (2) the major apex was excised, ‘major cut’, (3) the minor apex was excised, ‘minor cut’ or (4) both apices were excised ‘cut’ (Fig. 4D, E). The activity of angle meristems was subsequently monitored over 8 weeks (Fig. 4D, E). In ‘uncut’ plants, all angle meristems developed rhizophores following 5 weeks of growth, showing a similar pattern of development to the tissue culture system, and removal of either the major or the minor apex resulted in no conspicuous difference to ‘uncut’ plants (Fig. 4B, D, E). As in the tissue culture system, ‘cut’ plants with both the major and minor apices removed developed branches at a low frequency, implicating mobile signals from the shoot tips in angle meristem plasticity and suppression of branch outgrowth. However, unlike the tissue culture system (Fig. 4A-C), no rhizophore emergence followed decapitation in ‘cut’ plants in the whole plant system (Fig. 4B, E). This led us to infer that long-range cues from the base as well as the apex of the plant may regulate rhizophore development.

### The shoot apices are the main source of auxin

To investigate roles for apical auxin and long-range auxin transport in branch outgrowth from the angle meristems, we combined surgical experiments with a pharmacological approach applying NAA, NPA or a combination of NAA and NPA to excision sites (Fig. 4F, G). Following decapitation and 8 weeks of growth, plants with both tips excised and replaced with lanolin paste containing a solvent control produced branches (Fig. 4G). As in tissue culture (Fig. 4C), rhizophore development was restored in plants with both tips excised and replaced with a lanolin paste containing NAA (Fig. 4C, G) confirming that the apices act as an auxin source regulating angle meristem plasticity and branch outgrowth. However, a callus like tissue also initiated at a low frequency and rhizophores initiated at a lower frequency than in the tissue culture system (compare Fig. 4C with 4G). To test a role for auxin transport more directly, the auxin transport inhibitor NPA was applied in a lanolin paste to excision sites. Rhizophores and branches both initiated at low frequencies (Fig. 4G), and we considered that this may have been due to insufficient auxin production in remaining stem tissues to compensate for apex removal. Applying a lanolin paste with a combination of NAA and NPA did not restore rhizophore development to frequencies higher than the single NAA treatment but increased the frequency of production of the callus like tissue. Thus, NPA may have some local effects on auxin export from the angle meristems as well as affecting long-range transport from the shoot tips.

### *Three canonical* PIN*s are expressed in* S. kraussiana *shoots*

Using radiolabelled auxin transport assays in shoot cuttings, previously published work demonstrated that there is NPA-sensitive long-range basipetal auxin transport in *S. kraussiana* stems (Sanders and Langdale, 2013), and that such transport is mediated by PIN proteins in Arabidopsis (Gälweiler et al., 1998). To identify any potential involvement of PIN-mediated auxin transport in *S. kraussiana* bifurcation, previously identified *S. moellendorffii* PIN (SmPIN) peptide sequences were used in reciprocal tBLASTn searches against the *S. kraussiana* genome (Ge et al., 2016), enabling us to identify four *S. kraussiana PIN* (*SkPIN*) genes (Fig. 5A, B). Phylogenetic reconstruction identified *SkPINs* as orthologues of *SmPINR, SmPINS, SmPINT* and *SmPINV* respectively (Bennett, Brockington, et al., 2014), and they were named accordingly (Fig. 5B). The structure of each *SkPIN* (Fig. 5C) was determined by isolating and comparing cDNA sequences to genomic sequences using 3’ RACE, PCR, Sanger sequencing and sequence alignment. Whilst *SkPINR, SkPINS* and *SkPINT* had multiple introns towards the 3’ end of the gene, no introns were evident in *SkPINV* (Fig. 5C). To identify *PIN*s with potential roles in bifurcation, we used published RNASeq data (Ge et al., 2016) to generate heat maps that showed gross expression in shoot apices, stems, leaves, and rhizophores (Fig. 5D). Whilst *SkPINV* showed little to no expression, *SkPINR, SkPINS and SkPINT* were expressed in multiple shoot tissues. *SkPINR* and *SkPINS* were expressed in rhizophores, *SkPINS* and *SkPINT* were strongly expressed in shoot apices and *SkPINR, SkPINS* and *SkPINT* were strongly expressed in stems (Fig. 5D). Thus, we considered *SkPINR, SkPINS* and *SkPINT* candidate regulators of bifurcation.

**Figure 5:**
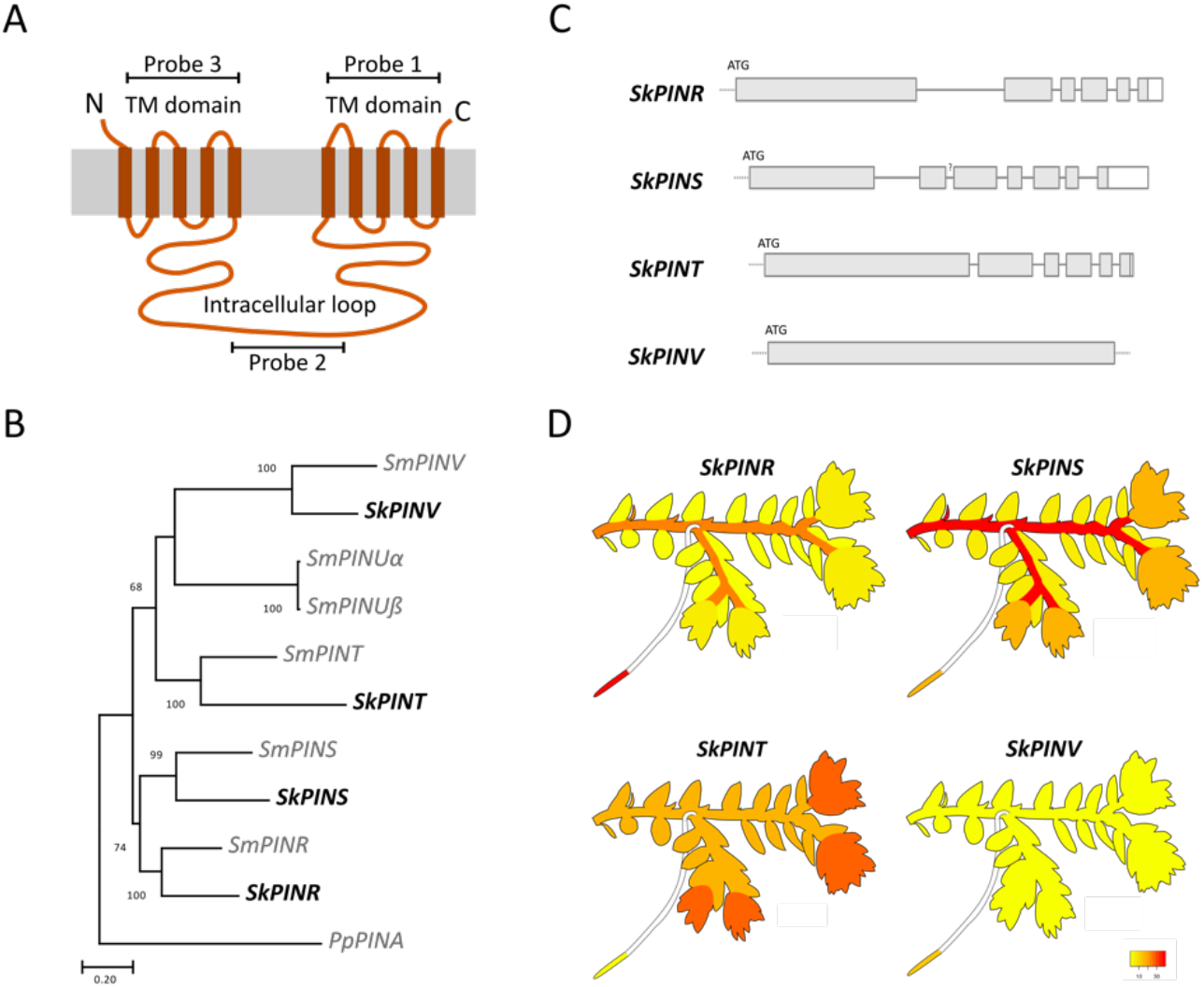
Three canonical PINs are expressed in *S. kraussiana* shoots. (**A**) Diagram showing the canonical PIN structure of lycophyte PINs characterised by a long intracellular loop and N- and C-terminal transmembrane domains. Positions of probes used for *in situ* hybridisation are shown. (**B**) Phylogenetic relationships between *S. moellendorffii* (*SmPIN*) and *S. kraussiana PIN* (*SkPIN*) genes. Maximum likelihood tree constructed using 226 amino acids of the N-terminal TM domain. Bootstrap values over 50 are shown. A canonical *Physcomitrium patens PIN* gene (*PpPINA*) was used to root the tree (Bennett, Brockington, et al., 2014). Gene names based on Bennett et al. (2014). No orthologs for *SmPINUα* or *SmPINUß* were detected, however *in silico* expression analyses suggest that *SmPINU* genes are not expressed in *S. moellendorffii* (Supplementary Fig. 5). (**C**) *S. kraussiana PIN* gene structures. Grey boxes represent exons, white boxes represent 5’/3’ UTRs and connecting lines represent introns. Dashed lines represent sequences that not been experimentally validated. (**D**) *In silico* expression analyses using RNA Seq data (Ge et al., 2016) showed tissue specific expression of *SkPINS* and *SkPINT* in the shoot apex, *SkPINR, SKPINS*, and *SkPINT* in mature stems, and *SkPINR* and *SkPINS* in rhizophore apices. *SkPINV* was not highly expressed in any tissue type. Heat maps represent relative expression levels with undetectable expression represented in yellow and the highest relative expression represented in red.

### SkPINR *and* SkPINS *are likely bifurcation regulators*

Data shown in Fig. 2 and Fig. 3 suggest that short-range auxin transport in *S. kraussiana* shoot apices regulates apical cell activity during bifurcation and that long-range auxin transport in the stem vasculature globally coordinates bifurcation. Hence we considered the shoot apices and stem vasculature as likely sites of *SkPIN* action. To better resolve spatiotemporal aspects of *SkPIN* activity, we used *in situ* hybridisation with probes against the transmembrane domains and intracellular loops (Fig. 5A) to determine gene expression patterns (Fig. 6, Supplementary Fig. 6). The apical cells and merophytes were most clearly visualized in sagittal sections (Fig. 6A) and *SkPINR* was expressed at low levels within and high levels directly beneath these cells, whilst *SkPINS* was expressed in the shoot tip but distally from the apical cells (Fig. 6A, B). Both genes were strongly expressed during leaf initiation and in the leaf vasculature at later stages of leaf development (Fig. 6A). *SkPINT* was expressed distally from the shoot tips in the vasculature (Fig. 6A, B), and we were unable to detect *SkPINV* expression in any tissue (Supplementary Fig. 6). Thus, *SkPINR* is the strongest candidate regulator of short-range auxin transport during bifurcation. Different stages of the bifurcation cycle were more clearly visualized in frontal sections (Fig. 6A, B). Again, these analyses showed *SkPINR* and *SKPINS* expression in shoot tips and initiating leaves, and they also showed *SkPINT* expression in developing ligules (Supplementary Fig. 6C). All three genes were expressed in the stem vasculature away from the shoot tips (Fig. 6A, B). As the vasculature is the likely site of long-range auxin transport, and *SkPINR* and *SkPINS* were most highly expressed in this tissue, we concluded that *SkPINR* and *SkPINS* likely act together to co-ordinate long-range auxin transport and overall patterns of bifurcation.

**Figure 6:**
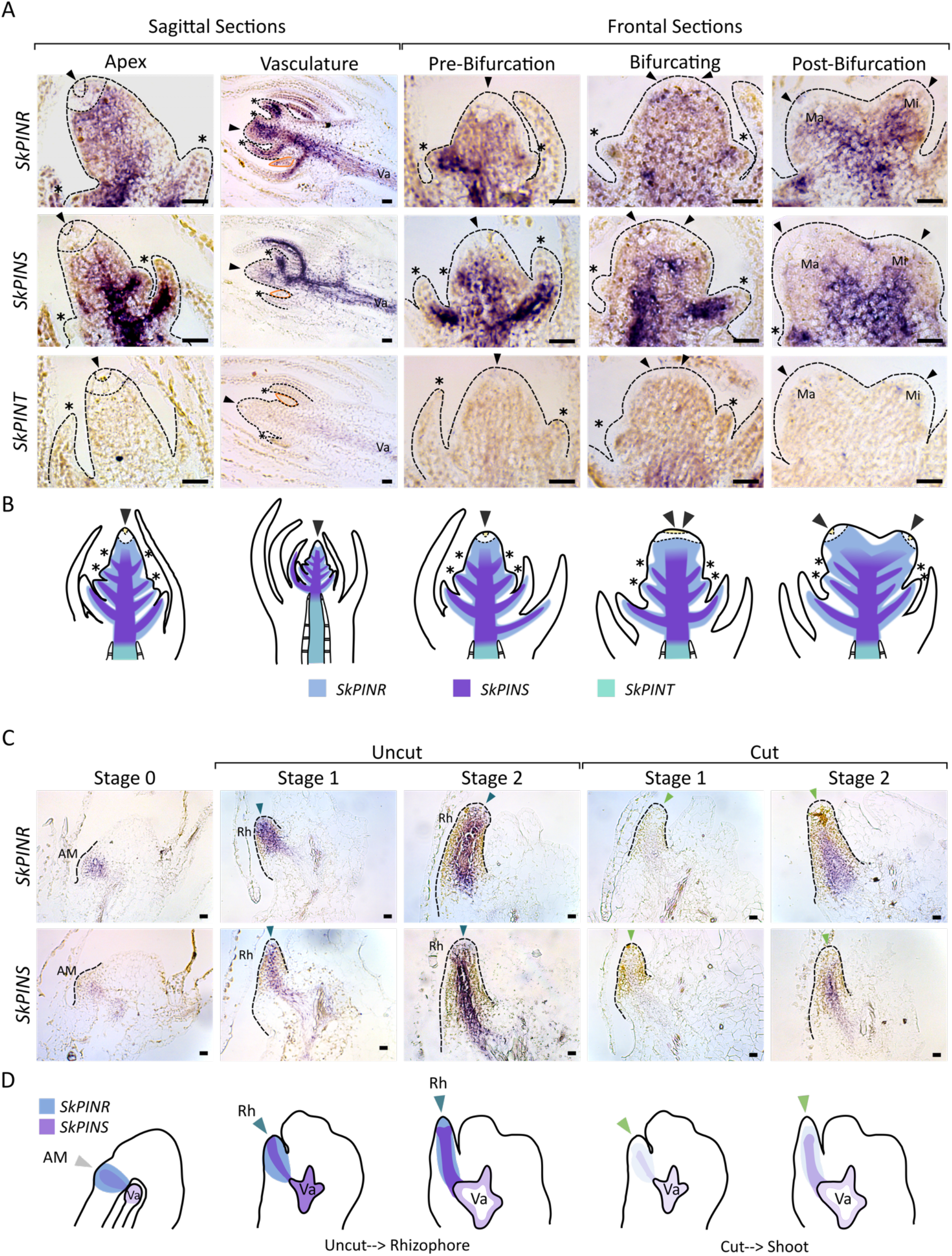
*SkPINR* and *SkPINS* are expressed in shoot apices, stem vasculature and rhizophore development. (**A**) Light micrographs of RNA *in situ* hybridization sections of the *S. kraussiana* apex during bifurcation. Dotted lines show the edge of the apex, the merophytes and the apical cell, and the Major (Ma) and Minor (Mi) branches are evident based on their relative size. Arrowheads = branch apex, asterisks = initiating leaves, orange outline = developing ligules. Va = vasculature. Scale bar = 0.02 mm. (**B**) Schematic summarising the expression patterns shown in **A**. *SkPINR* (blue) is broadly expressed across the shoot apex at all stages of bifurcation but is expressed to a lesser extent in the initial cells (yellow) and merophytes (dashed area). *SkPINS* (purple) has similar expression to *SkPINR* but is more closely associated with the developing vasculature. *SkPINT*(green) had no detectable apical expression but was expressed at later stages of vascular development. All three genes were expressed exclusively in the meristeles but not surrounding tissues. Asterisk = developing leaves, Black arrowhead = branch apex. (**C**) Light micrographs of RNA *in situ* hybridization sections of the angle meristem (stage 0) and developing rhizophores. Dotted lines show the edge of the angle meristem or rhizophore, blue arrowhead = rhizophore apex, green arrowhead = branch apex. Scale bar = 0.02 mm. (**D**) Schematics summarising the expression patterns shown in **C**. *SkPINR* (blue) is broadly expressed across the AM and rhizophore apex. *SkPINS* (purple) has similar expression to *SkPINR* but is more closely associated with the developing vasculature. One week after surgical decapitation, *SkPINR* and *SkPINS* were expressed less strongly at stages 1 and 2 of organ emergence than in uncut plants, likely implicating a decrease in *PIN* expression in the switch to branch identity.

### SkPINS *and* SkPINT *regulate angle meristem activity and plasticity*

As well as bifurcation, long-range auxin transport through the stem vasculature regulates angle meristem activity. To further investigate the potential involvement of *SkPINs* in branch outgrowth and angle meristem plasticity, we evaluated gene expression patterns at different stages of organ emergence from angle meristems in plants that were fixed with the tips intact or that were fixed one week following surgical decapitation (Fig. 6C, D). We found broad expression of *SkPINR* in angle meristems prior to organ emergence (Stage 0), but *SkPINS* was expressed less intensely than *SkPINR* (Fig. 6C, D). *SkPINT* and *SkPINV* expression was undetectable (Supplementary Fig. 6). *SkPINR* and *SkPINS* expression intensified and extended throughout the vasculature as rhizophores initiated (Stage 1) and emerged (Stage 2) from branch points where the shoot apices were intact (Fig. 6C, D). In contrast, *SkPINR* and *SkPINS* expression was barely detectable (Stage 1) or decreased relative to ‘uncut’ plants (Stage 2) during organ emergence in ‘cut’ plants (Fig. 6C, D). We therefore propose that a decrease in *SkPINR* and *SkPINS* expression may be involved in the switch from rhizophore to branch identity.

## Discussion

### PIN-mediated auxin transport globally co-ordinates bifurcation and angle meristem identity

We diagrammatically summarise our hypotheses relating to the evolution of branching and mechanisms underlying *S. kraussiana* branching in Fig. 7 (Fig. 7A-D). Overall, our results lead us to a model of bifurcation whereby short-range PIN-mediated auxin transport out of the apical cells is required to maintain their stem cell identity and promote bifurcation (Fig. 7B). The rate of bifurcation in major and minor branches is co-ordinated globally, and long-range auxin transport from different parts of the shoot system modulates branch dominance. The major branch tips are stronger auxin sources than the minor branch tips, and we propose that they have a greater capacity for auxin export enabling branch dominance (Fig. 7B). Surgical decapitation of the major branch tips enables minor branches to gain dominance (Fig. 7B). *SkPINR* is likely to provide the short-range auxin transport required to promote apical activity in bifurcation, and *SkPINR, SkPINS* and *SkPINT* are likely to act together to provide long-range auxin transport in the stem vasculature, globally coordinating bifurcation. Auxin export from the shoot tips also modulates the activity of angle meristems and *SkPINR* and *SkPINS* are likely to provide the long-range auxin transport required (Fig. 7C). In plants rooted on soil, an interruption to the flow of auxin by decapitation leads to a switch in angle meristem identity and the emergence of branches rather than rhizophores (Fig. 4E, Fig. 7C), and this switch is associated with a transitory drop in *SKPINR* and *SkPINS* expression levels (Fig. 6). However, in smaller explants grown in tissue culture, interrupting auxin flow by decapitation leads some angle meristems to develop branches, but the majority continue to develop rhizophores (Fig. 4B) suggesting that growth conditions or basal cues such as cytokinin or strigolactone (Domagalska and Leyser, 2011) may also regulate angle meristem activity and organ emergence. Taken together, our results suggest that PIN-mediated auxin transport is a critical regulator of the overall branching architecture of *S. kraussiana* (Fig. 7).

**Figure 7:**
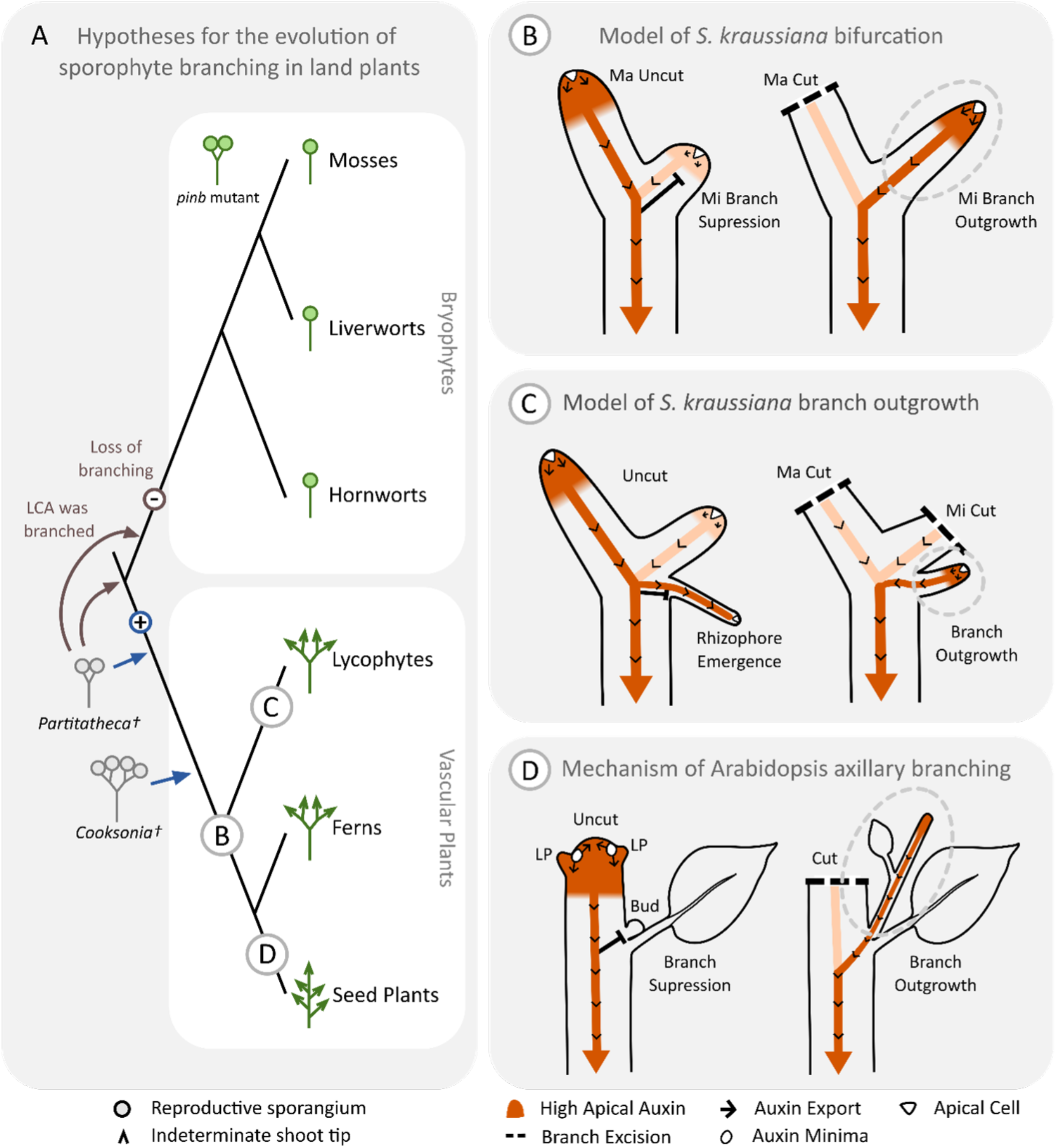
Hypotheses relating to mechanisms of bifurcation and plastic branch outgrowth in *Selaginella kraussiana* and to the evolution of branching mechanisms in land plants. **(A)** Current hypothesis of relationships between land plants showing the origin of bifurcation (+) and the independent evolution of plastic branch outgrowth in lycophytes and seed plants (C and D). Eophyte fossils such as *Partitatheca* and early vascular plant fossils such as *Cooksonia* have simple bifurcating forms with terminal sporangia (Edwards et al., 2021, 2022), and previous hypotheses (blue arrows) proposed that bifurcation was acquired in early steps of vascular plant evolution (Harrison 2017a; Harrison and Morris, 2018). Together with the finding that disruption of *pinb* function in *Physcomitrium patens* can lead to bifurcation (Bennett, Liu, et al., 2014), our finding that PIN-mediated auxin transport is an ancestral branch regulator within vascular plants suggests that the common ancestor of land plants may have had a branching form similar to *Partitatheca* (brown arrows), and that branching was lost during the evolution of bryophytes (-). Phylogeny redrawn from Puttick et al. (2018). (**B-D**) Models of branch regulation in *S. kraussiana* and *Arabidopsis*. The intensity of the orange shading represents the concentration of auxin, and arrows represent the direction of auxin transport. Black bars represent growth suppression and dashed lines indicates sites of apex excision. (**B**) In *S. kraussiana*, auxin transport out of the apical cells (white triangles) is required to maintain their identity and promote bifurcation. The branch tips are auxin sources and major branches produce and export more auxin through the stem vasculature than minor branches. Excision of the major branch tips interrupts basipetal auxin transport, enabling auxin from the minor branch tips to access the polar auxin transport stream, and releasing minor branch suppression. **(C)** Lycophytes in the Selaginellales innovated a unique organ system with plastic growth responses and the capacity to develop as a rhizophore or a branch (Banks, 2009; Spencer et al., 2021), and long-range auxin transport from the shoot tips regulates plasticity in *S. kraussiana*. If the shoot tips are intact, rhizophores establish an acropetal auxin transport stream and there is no branch outgrowth, but if the shoot tips are excised, the identity of the angle meristem changes leading to callus development or branch development. **(D)** In Arabidopsis, auxin export from the leaf axils (white ovals) enables axillary meristem establishment (Q. Wang et al., 2014; Y. Wang et al., 2014), and basipetal auxin transport from dominant apices blocks auxin export from axillary buds, suppressing branch outgrowth (Müller and Leyser, 2011). Following surgical excision of dominant apices, axillary buds activate auxin export leading to branch outgrowth (Thimann and Skoog, 1933; Prusinkiewicz et al., 2009). LP = leaf primordium.

### Auxin transport out of apical cells affects their activity and identity

Our model of bifurcation is consistent with previous findings that there is long-range auxin transport in the stem in *S. kraussiana* and that NPA treatment leads to apex termination (Sanders and Langdale, 2013). Although previous work noted no link between auxin transport and branching (Sanders and Langdale, 2013), we view this discrepancy as an artefact of different experimental strategies used. Sanders and Langdale (2013) grew explants in shaking liquid culture with NPA, and liquid immersion and the lack of a gravity vector could both affect branching. The regulation of branch outgrowth from angle meristems has been studied more extensively than bifurcation, and our results are consistent with previous findings made in other *Selaginella* species that apical auxin suppresses branch outgrowth (Williams, 1937a). Topical transport inhibition below the shoot apices in *S. willdenovii* (Wochok and Sussex, 1975) or on the underside of the stem in *S. moellendorffii* (Mello et al., 2019) led to branch outgrowth, consistent with the notion that polar auxin transport plays a role. Although basipetal transport in the stem tissues enables rhizophore emergence, transport is acropetal in rhizophores, and this redirection of auxin transport is reflected in patterns of vascular development in many species of *Selaginella* (Matsunaga et al., 2017). Our own experiments attempting to disrupt basipetal transport by applying auxin transport inhibitors around the stem (500 μM NPA in lanolin) yielded no difference in angle meristem identity from controls, we think due to poor tissue penetration (data not shown). Our model of branch outgrowth from the angle meristems is that following apical decapitation, a loss in the strength of the auxin transport stream in the stems results in a drop in *SkPINR* and *SkPINS* expression in the angle meristems permitting changes in apical cell identity (Fig. 7). We hypothesise that if angle meristem apical cells accumulate auxin they proliferate to form callus or gain rhizophore identity, but if they can export sufficient auxin, branch outgrowth activates (Fig. 7C). Thus, the level of auxin transport out of apical cells may determine their identity.

### PIN-mediated auxin transport was an ancestral regulator of branching co-opted independently into lycophyte and seed plant plasticity

In conjunction with findings from *Physcomitrium patens* that PIN-mediated auxin transport is required to maintain apical cell identity and that disruption of PIN function can induce bifurcation (Bennett, Liu, et al., 2014), our data suggest that PIN-mediated auxin transport may be an ancestral requirement of land plant stem cell function and an ancestral regulator of branching within vascular plants (Fig. 7A). As some of the earliest land plant macrofossils comprise bifurcating axes with terminal sporangia but no specialised water conducting cells (e.g. Edwards et al., 2021, 2022), and there is growing evidence of evolutionary loss in the bryophytes (Harris et al., 2020, 2021), we speculate that the single-stemmed state of bryophyte sporophytes was derived from a last common ancestor of land plants manifesting PIN-regulated bifurcation (Fig. 7A). PINs underwent independent duplications in lycophytes and seed plants, and they were independently co-opted to regulate the plastic development of a unique organ system in *Selaginella* and axillary branching in seed plants during evolution (Fig. 7A, D). More broadly, many plant groups have evolved multicellular forms with a branching habit and apical dominance (Harrison, 2017a; Coudert et al., 2019). In conjunction with findings presented here, the recent report that *PpPINA, PpPINB* and *PpPINC* regulate apical dominance in moss filament branching (Nemec-Venza et al., 2022) suggests PIN-mediated auxin transport as a core mechanism for branching within land plants. We propose that heterotopic changes in PIN expression or localisation were key drivers of the evolution of diverse branching forms.

## Methods

### Plant growth conditions

*Selaginella kraussiana* (Kunze) A. Braun plants were grown in indirect light at 5-20 μmol m^-2^ sec^-1^, 21 °C, 70 % humidity in long day conditions (16 hrs light, 8 hrs dark). Plants were propagated by transferring apical shoots with at least three major axis bifurcations to soil (Levington Advance Pot and Bedding Compost) and grown in trays sealed in bags to increase humidity. Alternatively, explants were grown in axenic tissue culture with ¼ Gamborg B5 medium (Duchefa Biochemie; G0209.0050), 0.8 % Agar (Sigma; A4675-1KG), pH 5.8 and 1 % Sucrose (Alfa Aesar; A15583), in the same light conditions. To sterilise tissue, *S. kraussiana* explants with ~ 2 bifurcations on the major axis were harvested and rinsed with dH2O for 15 minutes. Explants were then incubated in 20 % Sodium Hypochlorite Solution (Fisher Scientific; S/5040/PB17) for 10 minutes, during which the falcon tube would be inverted 6 times every minute. Tissue was then rinsed 4 times in sterile dH2O before transfer to media. Sterile tissue was propagated in magenta boxes and hormone experiments were performed in deep petri dishes of varying sizes.

### Pharmacological experiments

Sterile explants with 1 bifurcation point were transferred to deep petri dishes containing ¼ strength Gamborg B5 media and 0.8 % Agar, at pH 5.8 with 1 % Sucrose (unless stated otherwise). Hormone treatments were performed at concentrations described in the results section. In all cases, solvents were standardised to 0.7 % Ethanol and 0.02% DMSO. Plants were grown for 8 weeks and photographed for phenotypic analysis. Angle meristem activity was observed once every week to check for the presence or absence of shoots and roots over a period of 5 weeks.

### Surgical experiments

To investigate bifurcation, explants with three bifurcations of the major axis were transferred to pots and grown in bags. Surgical interventions were made at week 0, and at week 8 plants were scanned using a HP Scanjet G2710 and the number of bifurcations at each side branch recorded. Using the same set up, surgical experiments were combined with pharmacological treatments described in the results section. Solvents were standardised to 7 % Ethanol and 0.2 % DMSO in all experiments. To investigate AM activity, explants with four bifurcations on the major axis were transferred to pots as above and grown for four weeks. The most distal shoot tips were decapitated and branches beneath the tips were marked with coloured beads. Plants were either left to grow (“Uncut”) or had the shoot tips excised (“Both Cut”). Lanolin hormone paste was applied with a syringe to excision points at concentrations stipulated in the results section. AM activity was observed weekly and the presence of a shoot branch or rhizophore recorded.

### *PIN* gene identification in *Selaginella kraussiana*

*Selaginella moellendorffii* PIN peptides were used in tBLASTn searches against the *Selaginella kraussiana* genome downloaded from Ge et al. (2016). An e-value cut off of 0.05 was used. The *S. kraussiana* sequence boundaries returned did not fall within annotated regions of the genome, so contigs to which the *S. moellendorffii* gene aligned were downloaded. Each contig was aligned to *S. moellendorffii* PIN genes to find regions of homology likely to encode SkPIN orthologues. Each putative *S. kraussiana* PIN orthologue was translated and aligned to *S. moellendorffii* orthologues using Clustal X (Thompson et al., 1997). Maximum Likelihood (ML) Trees were constructed in MEGA X (Kumar et al., 2018), using 226 conserved amino acids in exon 1 with the JTT Model of amino acid substitution (Jones et al., 1992), and including the equivalent sequence from *Physcomitrium patens PIN* (*PINA*) as an outgroup. The phylogenetic trees generated were used to name *S. kraussiana PIN* genes relative to their closest *S. moellendorffii* orthologue (Bennett, Brockington, et al., 2014). 4 *PIN* genes (*PINR, PINS, PINT*, and *PINV*) were identified.

### RNA Seq Gene expression quantification

*PIN* gene sequences were used in BLAST searches against the *S. kraussiana* genome assembly (Ge et al., 2016), to produce contig coordinates for each gene and create a simplified annotation format (SAF) file. Using these newly created SAF files and bam files kindly provided by the authors of Ge et al. (2016) for each tissue type (stem, leaves, root, shoot apex), the number of reads aligning to specific regions was determined using Featurecounts (Liao et al., 2014). Multimapping was allowed as total counts from multimapping did not differ dramatically from single mapping counts, but did allow quantification of *SkPINV*, as this gene has three identical sequences on three different contigs. Multimapping counts were used for all genes and normalised using FPKM. Heat map colours were generated with these values using http://www.heatmapper.ca/expression/ and overlayed onto *S. kraussiana* diagrams in Inkscape.

### RNA extraction

A modified Trizol prep was used to extract RNA (Chomczynski and Sacchi, 1987). 500 μL of Solution D (4M guanidine isothiocyanate, 25 mM sodium citrate pH 7.0, 0.5 % sarkosyl), 500 μl Phenol-Chloroform-Isoamyl alcohol mixture (Sigma; 77618) and 50 μl 2M sodium acetate pH 4 were mixed and added to 30 mg of frozen, ground, *S. kraussiana* shoot tips. The solution was vortexed for 10 seconds and the supernatant transferred to a new tube. After 5 minutes at room temperature, 200 μl Chloroform: Isoamyl alcohol 24:1 (Serva; 39554.02) was added and incubated for 5 minutes. The samples were centrifuged at 12,000 x g for 15 minutes at 4 °C. 500 μl of the top aqueous layer was added to 500 μl isopropanol. After 10 minutes incubation at room temperature, solutions were mixed by inverting, then centrifuged at 12, 000 x g for 10 minutes at 4 °C. The white RNA pellet was washed in 100 % Ethanol, centrifuged again at 7,500 g for 4 minutes at 4 °C, air dried, then resuspended in 30 μl DEPC H20. RNA quality was checked by gel electrophoresis and concentrations were determined using a NanoPhotometer^®^ N60 (Implen).

### Primers

A list of primers for PCR and cloning is included in Table 1.

**Table 1.**
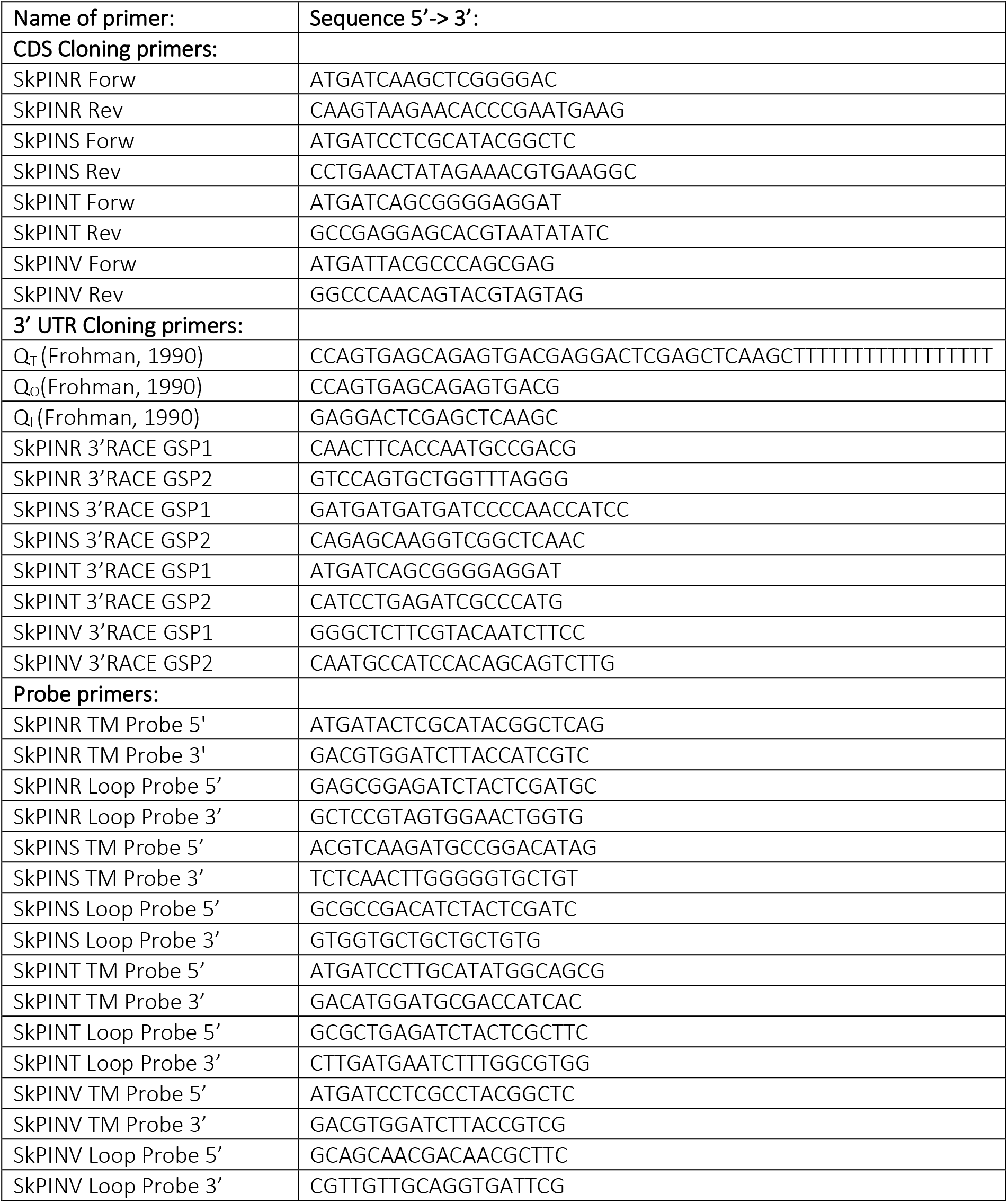
Primers used in this study.

### cDNA synthesis

1 μg of *S. kraussiana* RNA was incubated with DNAse I (Thermo Scientific; EN0521) and 1 x Reaction Buffer in a total volume of 10 μl for 30 minutes at 37 °C. 1 μl of 50 mM EDTA was then added and the solution incubated at 65 °C for 10 minutes. This whole reaction was added to Primer Mix (Qiagen; 1030542), 0.5 mM dNTPs, 1 x First Strand Buffer, 0.01 M DTT, RiboLock RNAse Inhibitor (Thermo Scientific; E0038) and Superscript^®^ II Reverse Transcriptase (Invitrogen; 100004925) to a final volume of 20 μl. The reaction was incubated at 42 °C for 50 minutes and then 70 °C for 15 minutes. A no-RT control sample was also produced for every round of cDNA synthesis to check for contamination.

### PIN CDS cloning

Primers to amplify *S. kraussiana PIN* genes were designed by aligning *S. moellendorffii* CDS to the homologous *S. kraussiana* genomic sequence. Primers were located at regions of high sequence similarity and as close as possible to the predicted ATG and stop codon of the transcript. All primers were checked for low self-dimerization and heterodimerisation ΔG values using IDT OligoAnalyser software (https://eu.idtdna.com/pages/tools/oligoanalyzer?returnurl=%2Fcalc%2Fanalyzer). PCR was performed with *S. kraussiana* cDNA and Q5^®^ HF DNA Polymerase (NEB Biolabs; M0491L). PCR products were separated by gel electrophoresis and extracted, incubated with Taq Polymerase for 30 minutes at 68 °C for A-tail addition, and ligated into pGEM^®^-T Easy Vector (Promega; A137A) overnight at 16 °C according to the manufacturer’s protocol. After transformation into TOP10 *E. coli* by electroporation, positive colonies were screened by digest or culture PCR, and then sequenced by Eurofins Genomics TubeSeq Service. Sequence results were aligned to the *S. kraussiana* genome, to identify exon and intron positions.

### 3’ RACE PCR

To identify the 3’ UTR of SkPINs, 3’ RACE PCR was performed. cDNA was synthesised as described above, but using the Q_T_ primer (Frohman, 1990). The first round of PCR used a Forward Gene Specific Primer (GSP1) within a known exon in the middle of the transcript, and the Q_O_ primer (Frohman, 1990). The second round of PCR used the PCR product from the first round, a Forward Gene Specific Primer (GSP2) nested further downstream of GSP1, and the Q_I_ primer (Frohman, 1990). Q5 HF DNA Polymerase was used for all reactions according to the manufacturer’s guidelines. For the 1^st^ PCR, a 2 minute extension with 58 °C annealing temperature and 25 cycles were used. For the 2^nd^ PCR, 35 cycles were used. The PCR product from the 2^nd^ PCR was visualized by gel electrophoresis and extracted, incubated with Taq polymerase for 30 minutes at 68 °C, and ligated into pGEM-T Easy overnight at 16 °C according to the manufacturer’s protocol. After transformation into TOP10 *E. coli* by electroporation, positive colonies were screened by digest or culture PCR, and then sequenced by Eurofins Genomics TubeSeq Service. Sequence results were aligned to the *S. kraussiana* genome to identify the 3’ UTR of each gene.

### RNA probe synthesis

pGEM-T-Easy DNA vectors containing ~300bp of each *S. kraussiana PIN* gene in both orientations were amplified using a midiprep kit (Qiagen; 12143). Plasmids were digested with SpeI (NEB Biolabs; R3133S) and purified by phenol/chloroform extraction. RNA transcription in sense and antisense orientations was performed with the T7 RNA polymerase (NEB Biolabs; M0251S) using Digoxigenin-11-UTP (Roche; 11209256910) to label the probes. After DNAse treatment, RNA was precipitated with 7.5 M NH4Ac and cold ethanol before probe hydrolysis (time depended on length of template). Probe fragments were then precipitated with 3 M NaOAc, 10 % HAc and ethanol, and re-dissolved in a 50:50 mix of DEPC treated H20 and formamide. To verify DIG labelling efficiency, probe dilutions were spotted onto a Bright Star Plus Nylon membrane (Invitrogen, AM10102), washed for 5 mins in Buffer 1 (100 mM Tris-HCl, 150 mM NaCl), blocked for 30 mins with buffer 2 (0.5 % (w/v) blocking reagent [Roche, 11096176001] in buffer 1), washed 3 times in buffer 1 for 5 mins, and incubated for 30 mins in 1:5000 Anti-Digoxigenin- AP Fab Fragment antibodies (Roche; 11093274910). After washing the membrane twice in buffer 1 for 15 mins, the signal was developed using a NBT/ BCIP solution pH 9.5 [Sigma-Aldrich, B5655].

### Tissue fixation for *in situ* hybridisation

Shoot apices and bifurcation points were dissected and fixed in 4 % Paraformaldehyde (Sigma-Aldrich, 441244) + 4 % DMSO overnight at 4 °C. Tissue was passed through a 4 °C ethanol series (30 %, 40 %, 50 %, 60 %, 70 %, for an hour each) before transfer to a Tissue-Tek VIP (Sakura). Tissue was processed as follows: 70 % ethanol (1 hr), 80 % ethanol (1 hr), 90 % ethanol (1 hr), 95 % ethanol (1 hr), 100 % ethanol (1 hr), 100 % ethanol (1 hr), 100 % ethanol (1.5 hr), 100 % Histoclear (National Diagnostics; A2-0101) (1 hr), 100 % Histoclear (1 hr), 100 % Histoclear (1.5 hr). All of these steps were performed at 35 °C with a slow mix. Tissue was transferred to fresh wax for 1.5 hrs, 2 hrs, 2.5 hrs, 2.5 hrs, all at 60 °C. Tissue was embedded in moulds and stored at 4 °C until use. 8 μm sections were prepared using a Leica RM2245 microtome and left overnight on a 42 °C hotplate before use.

### *In situ* hybridisation

Sections were pre-treated as follows: Histoclear (10 mins), Histoclear (10 mins), 100 % ethanol (1 min), 100 % ethanol (30 s), 95 % ethanol (30 s), 85 % ethanol + 0.85 % saline (30 s), 50 % ethanol + 0.85 % saline (30 s), 30 % ethanol + 0.85 % saline (30 s), 0.85 % saline (2 mins), Phosphate Buffered Saline (PBS) (2 mins), 0.125 mg ml^-1^ Pronase (37°C, 10 mins) (Sigma-Aldrich, P2308), 0.2 % Glycine (2 mins) (ChemCruz, sc-29096), PBS (2 mins), 4 % Paraformaldehyde in PBS (10 mins), PBS (2 mins), PBS (2 mins), Acetic anhydride (Sigma-Aldrich, 320102) in 0.1 M triethanolamine pH 8 (10 mins) (Sigma-Aldrich, T58300), PBS (2 min), 0.85 % saline (2 mins), 30 % ethanol + 0.85 % saline (30 secs), 50 % ethanol + 0.85 % saline (30 secs), 85 % ethanol + 0.85 % saline (30 secs), 95 % ethanol (30 sec), 100 % ethanol (30 sec), fresh 100 % ethanol (30 sec). Slides were left to dry for 1-2 hours.

Probes were mixed in a 1:4 ratio with hybridisation buffer (300 mM NaCl, 10 mM Tris-HCl pH 6.8, 10 mM NaPO4 buffer, 5 mM EDTA, 50 % deionised formamide, 1 mg ml^-1^ tRNA, 1 x Denhardt’s solution [Sigma-Aldrich, D2532], 10 % Dextran Sulphate [Sigma-Aldrich, D8906]) and applied to slides. Hybridisations were incubated at 50 °C overnight, before washing for 1 hour 30 mins at 50 °C in wash solution (15 mM NaCl, 1.5 mM Na3C6H5O7). Following two incubations in NTE solution (0.5 M NaCl, 10 mM Tris-HCl pH 7.5, 1 mM EDTA) for 5 mins at 37 °C, slides were transferred to NTE + 20 μg/ml RNAse A for 30 mins at 37 °C. Slides were then washed in NTE x 3 (5 mins each, 37 °C), wash solution (1 hr, 50 °C), and PBS (5 mins).

Anti-DIG antibodies were applied and signal developed as follows: 5 mins in Buffer 1 (100 mM Tris-HCl, 150 mM NaCl), 30 mins in Buffer 2 (0.5 % (w/v) blocking reagent in buffer 1), 30 mins in Buffer 3 (1 % BSA, 0.3 % Triton X-100 in Buffer 1), 1.5 hours in Buffer 4 (Anti-digoxigenin-AP 1:3000 in Buffer 3), 4 x 20 mins in Buffer 3, 5 mins in Buffer 1, 5 mins in Buffer 5 (100 mM Tris pH 9.5, 100 mM NaCl, 50 mM MgCl2), followed by 12 hours+ in Buffer 6 (NBT/ BCIP solution) until the signal develops. The enzymatic reaction was then stopped by washing slides for 30 sec each in distilled H20, 70 % ethanol, 95 % ethanol, 100 % ethanol, 95 % ethanol, 70 % ethanol, and distilled H20. Sections were mounted in Entellan mounting medium (Merck, 1.07961.0500) with coverslips before imaging.

### Image acquisition

Images of living *S. kraussiana* were taken using a Keyence Digital microscope (VHX-1000E) with a RZ x 50 or RX x 20-x 200 objective lens, a D80 Nikon camera with a EX Sigma 50mm 1:2.8 DG MACRO lens, or a Google Pixel 6 phone camera. Images of *in situ* sections were taken using a Leica DM2000 LED microscope with x 20 and x 40 objective lenses and a Leica MC120 HD camera attachment.

### Data analysis

All data were collected and bar graphs produced in Microsoft Excel (2020). Any statistical analysis and boxplots were generated in R studio (RStudio Team (2020)). RStudio: Integrated Development for R. RStudio, PBC, Boston, MA URL http://www.rstudio.com/). All figures and diagrams were produced using Inkscape (Inkscape project (2020) https://inkscape.org/.).

## Acknowledgements

We thank Jane Langdale and Julie Bull for providing plant tissue and use of histology facilities the authors of Ge et al., 2016 for sharing their data, and Alex Paterson and Ben White for assistance with bioinformatic analyses. We thank members of the Harrison Lab for comments on a manuscript draft. We thank the Leverhulme Trust for funding our work (RPG-2018-220).

## Author contributions

C. J.H. conceived the project. C.J.H, V.S. and L.B. designed the experiments. V.S. did the experimental work, analysed data and prepared all the figures except Figure 1D (C.J.H). L.B. assisted with preliminary decapitation experiments. V.S. and C.J.H. wrote the manuscript.

---

**Supplementary Figure 1 related to Figure 1:**
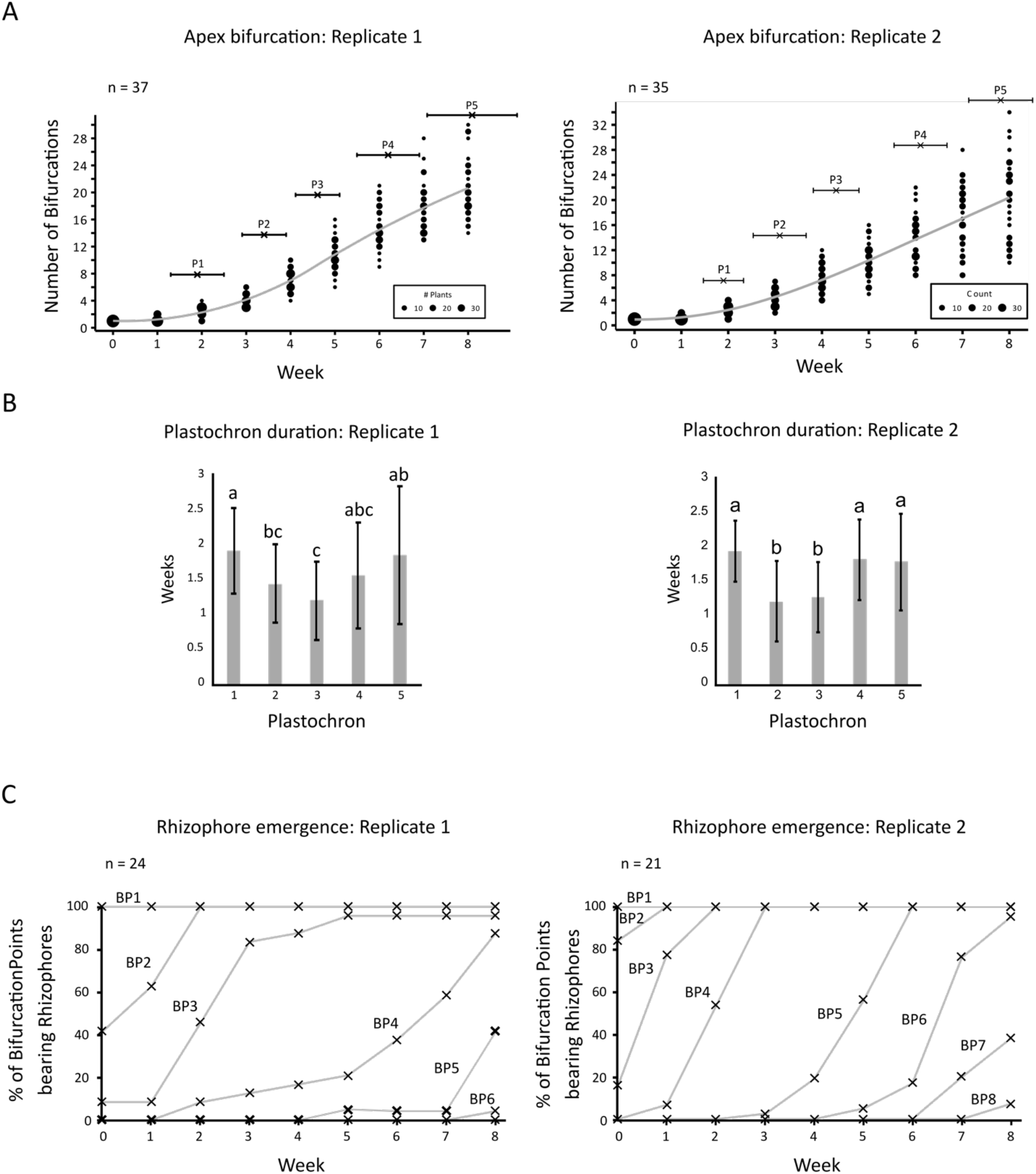
Bifurcation and rhizophore emergence time course data from two experimental replicates. (A) Graphs showing the number of bifurcations of the main apex following explant transfer to soil. (B) Bar graphs showing the mean duration of plastochrons 1-5. P2 and P3 proceeded more rapidly than P1, P4 and P5. One-way ANOVAs with Kruskal-Wallis tests for multiple comparisons were performed (χ^2^(4)= 22.87, p=0.00014; χ^2^(4)= 42.07, p<0.0001). Error bars show standard deviation. (C) Graphs showing the percentage of bifurcation points (BPs) bearing a rhizophore in an eight-week time course.

**Supplementary Figure 2 related to Figure 2:**
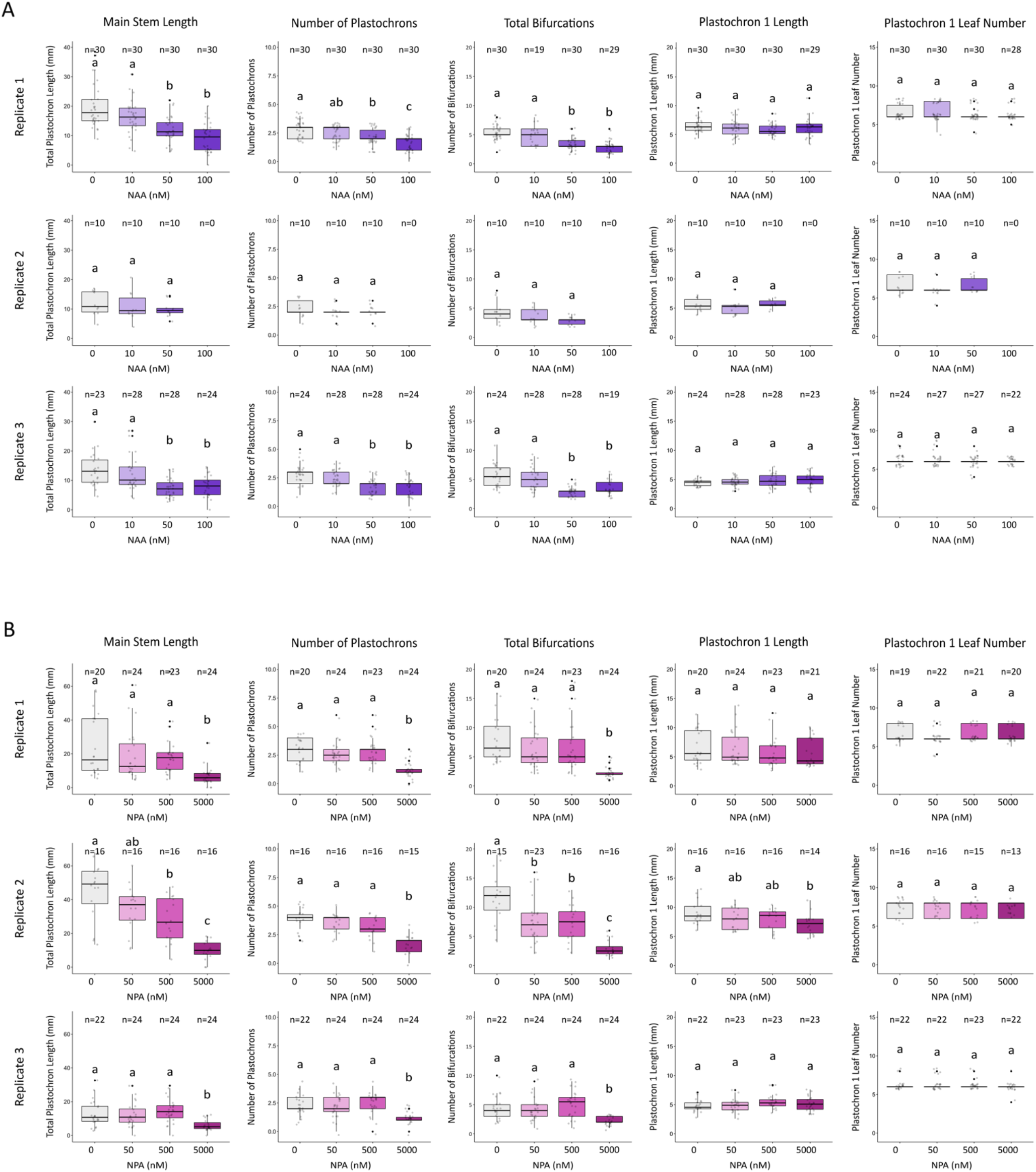
Auxin and auxin transport inhibitor application suppress bifurcation. (**A**) Graphs representing data from three experimental replicates of pharmacological experiments using explants with one branch point and grown in tissue culture for eight weeks with 0 nM, 10 nM, 50 nM or 100 nM NAA. (**B**) Graphs representing data from three experimental replicates of pharmacological experiments using explants with one branch point and grown in tissue culture for eight weeks with 0 nM, 50 nM, 500 nM or 5 μM NPA. In all graphs, boxes represent lower quartile, median and upper quartile. Lines represent spread of data and black points show outliers. One-way ANOVA tests with Tukey or Kruskal Wallis multiple comparisons were performed. (NAA Rep 1: F(3,116)= 20.44, p<0.0001; F(3,116)= 18.9, p<0.0001; F(3,104)= 26.21, p<0.0001; F(3,115)= 1.33, p=0.27; χ^2^(3)= 2.01, p=0.57. NAA Rep 2: F(2,27)= 0.58, p=0.57; F(2,27)= 0.36, p=0.7; F(2,27)= 2.94, p=0.07; F(2,27)= 0.66, p=0.53; χ^2^(2)= 2.15, p=0.34. NAA Rep 3: F(3,99)= 10.79, p<0.0001; F(3,100)= 12.26, p<0.0001; F(3,95)= 18.2, p<0.0001; F(3,99)= 1.37, p=0.26; χ^2^(3)=6.13, p=0.1. NPA Rep 1: F(3,87)= 7.07, p=0.00026; F(3,87)= 16.6, p<0.0001; F(3,87)= 12.22, p<0.0001; F(3,84)= 0.59, p=0.62; χ^2^(3)= 3.05, p=0.38. NPA Rep 2: F(3,60)= 19.94, p<0.0001; F(3,59)= 17.44, p<0.0001; F(3,66)= 17.94, p<0.0001; F(3,58)= 2.45, p=0.072; χ^2^(3)= 0.42, p=0.94. NPA Rep 3: F(3,90)= 7.48, p=0.00016; F(3,90)= 9.79, p<0.0001; F(3,90)= 11.86, p<0.0001; F(3,87)= 1.89, p=0.14; χ^2^(3)= 4.3, p=0.23).

**Supplementary Figure 3 related to Figure 3:**
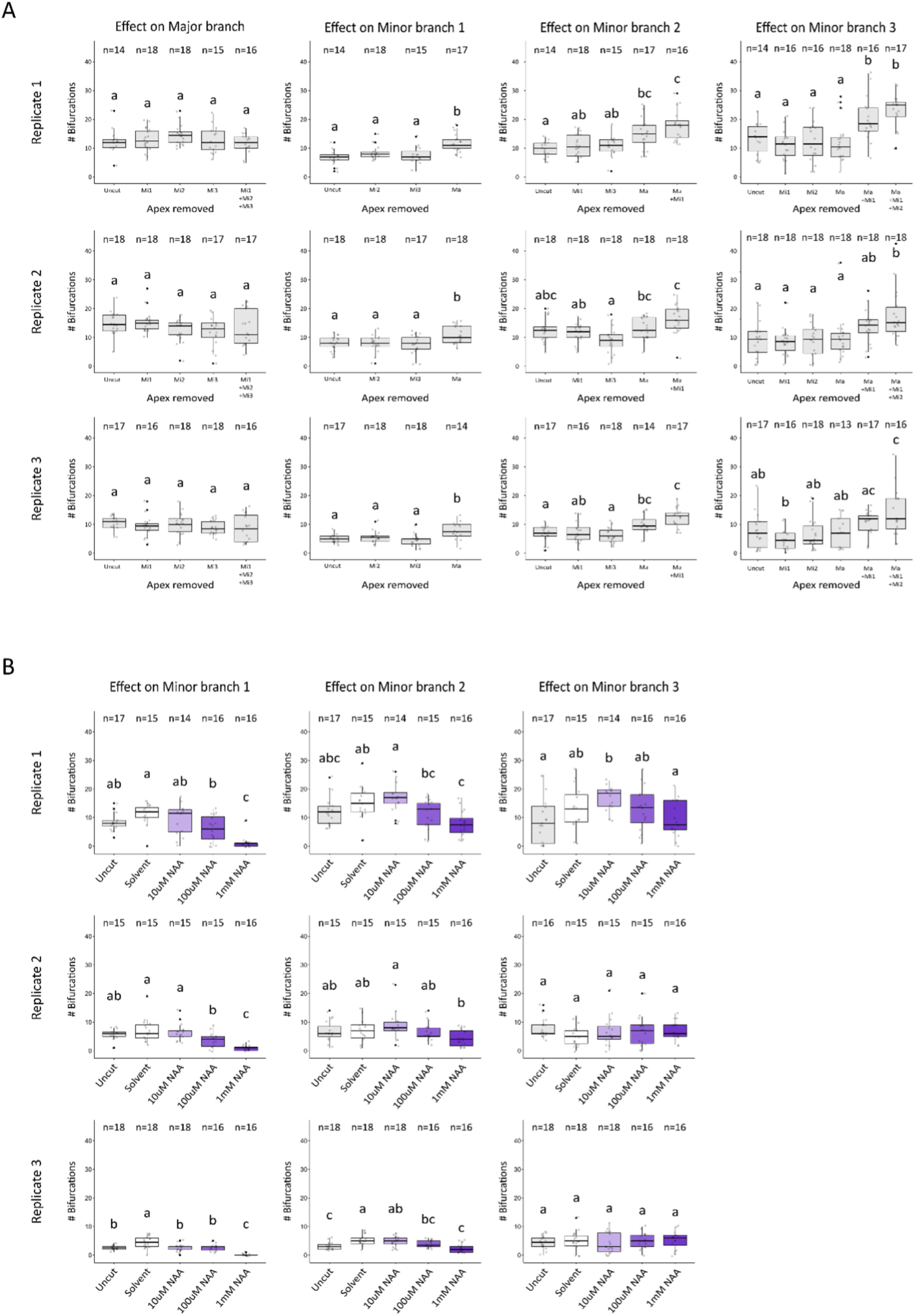
Data from apex decapitation and lanolin-auxin application replicates. (**A**) Graphs representing data from three experimental replicates of decapitation experiments using explants with three branch points and grown in soil for eight weeks. Either the major, minor 1, minor 2, minor 3 or a combination of these branches were removed, and the subsequent bifurcation rates recorded. In all graphs, boxes represent lower quartile, median and upper quartile. Lines represent spread of data and black points show outliers. Oneway ANOVAs with Tukey tests for multiple comparisons were performed (F(4,76)= 1.28, p=0.28; F(3,60)= 9.83, p<0.0001; F(4,75)= 9.94, p<0.0001; F(5,91)= 8.95, p<0.0001; F(4,83)= 1.73, p=0.15; F(3,67)= 5.39, p=0.002; F(4,85)= 6.7, p<0.0001; F(5,102)= 5.19, p=0.00028; F(4,80)= 0.99, p=0.42; F(3,63)= 6.62, p=0.00059; F(4,77)= 13.2, p<0.0001; F(5,91)= 6.08, p<0.0001). (**B**) Graphs showing data from three experimental replicates for replacement of the major branch with lanolin impregnated with varying concentrations of auxin. Explants were grown for eight weeks on soil. In all graphs, boxes represent lower quartile, median and upper quartile. Lines represent spread of data and black points show outliers. One-way ANOVAs with Tukey tests for multiple comparisons were performed (F(4,73)= 14.63, p<0.0001; F(4,72)= 6.86, p<0.0001; F(4,73)= 2.91, p=0.027; F(4,71)= 12.95, p<0.0001; F(4,71)= 2.76, p=0.034; F(4,72)= 0.81, p=0.52; F(4,81)= 24.77, p<0.0001; F(4,81)= 10.54, p<0.0001; F(4,81)= 0.35, p=0.84).

**Supplementary Figure 4 related to Figure 4:**
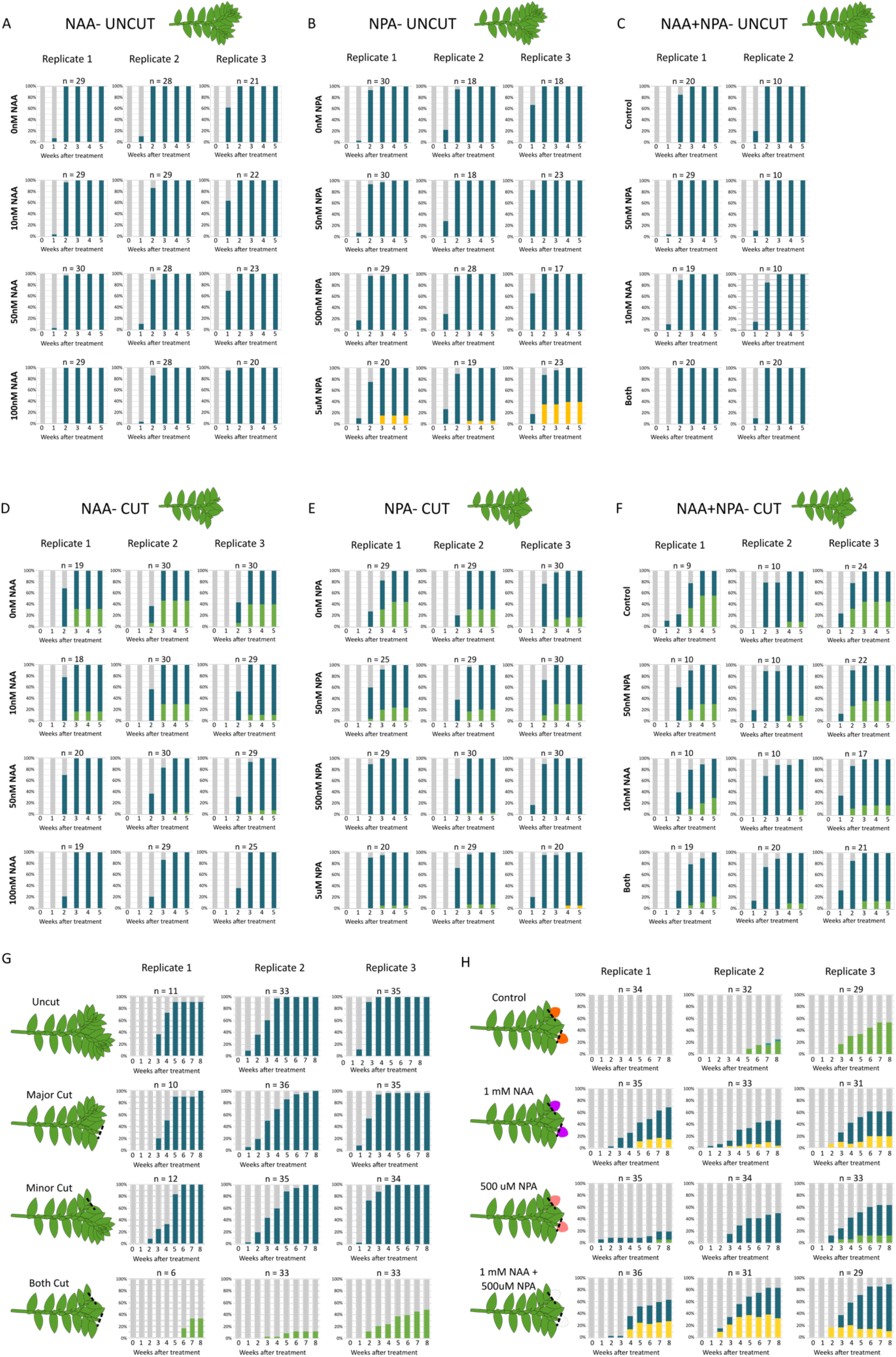
Effect of surgical decapitations and pharmacological treatments on rhizophore emergence. (**A, D**) Graphs representing data from three experimental replicates grown in axenic culture with 0 nM, 10 nM, 50 nM or 100 nM NAA. Explants had 1 BP and the apices were (**A**) left intact (Uncut) or (**D**) removed (Cut). (**B, E**) Graphs representing data from three experimental replicates grown in axenic culture with 0 nM, 50 nM, 500 nM or 5 μM NPA. Explants had 1 BP and the apices were (**B**) left intact (Uncut) or (**E**) removed (Cut). (**C, F**) Graphs representing data from two/ three experimental replicates grown in axenic culture with 0 nM, 10 nM NAA, 50 nM NPA or 10 nM NAA + 50 nM NPA. Explants had 1 BP and the apices were (**C**) left intact (Uncut) or (**F**) removed (Cut). (**G**) Graphs representing data from three replicates of decapitation experiments. (**H**) Graphs representing data from three replicates of experiments using decapitation and replacement of decapitated apices with lanolin paste containing 1 mM NAA, 500 μM NPA or a combination of 1 mM NAA and 500 μM NPA. In all graphs, grey shading indicates angle meristem identity, blue shading indicates rhizophore identity, green shading indicates branch identity and yellow shading indicates callus.

**Supplementary Figure 5 related to Figure 5:**
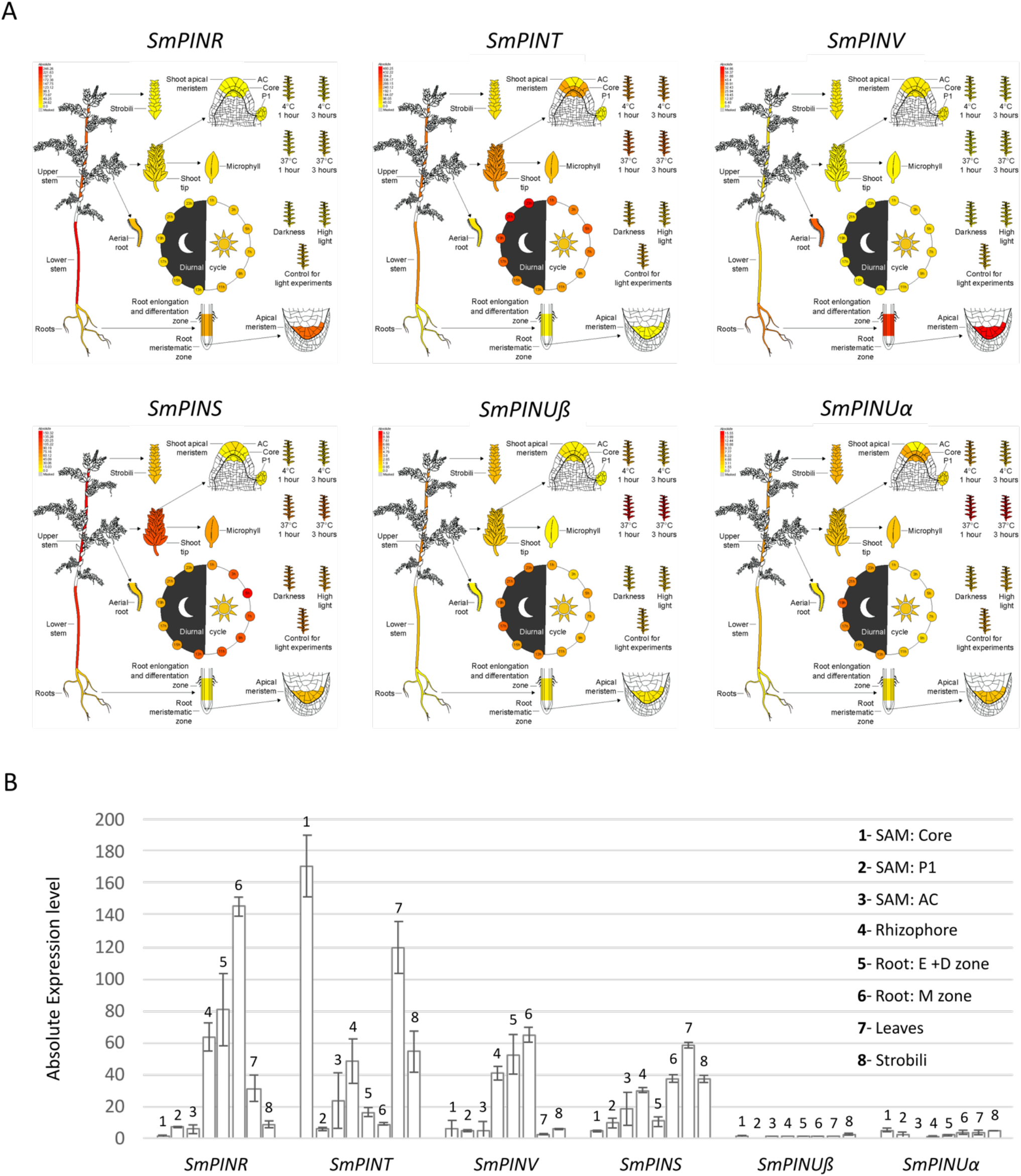
Selaginella moellendorffii *PIN in silico* expression data. (**A-B**) *In silico* RNA-Seq expression analyses of *S. moellendorffii PIN* genes from the BAR *Selaginella* eFP browser (Ferrari et al., 2020). The same data are represented as heat maps (**A**) and bar graphs (**B**). *SmPINR, SmPINT, SmPINV* and *SmPINS* are expressed variably across different tissues. *SmPINUα* and *SmPINUß* are not expressed as highly in any tissue. *SmPINR* is upregulated in the maturation zone of the zoot; *SmPINT* is upregulated in the SAM core; *SmPINV* is upregulated in all regions of the root; and *SmPINS* is upregulated in the leaves.

**Supplementary Figure 6 related to Figure 6:**
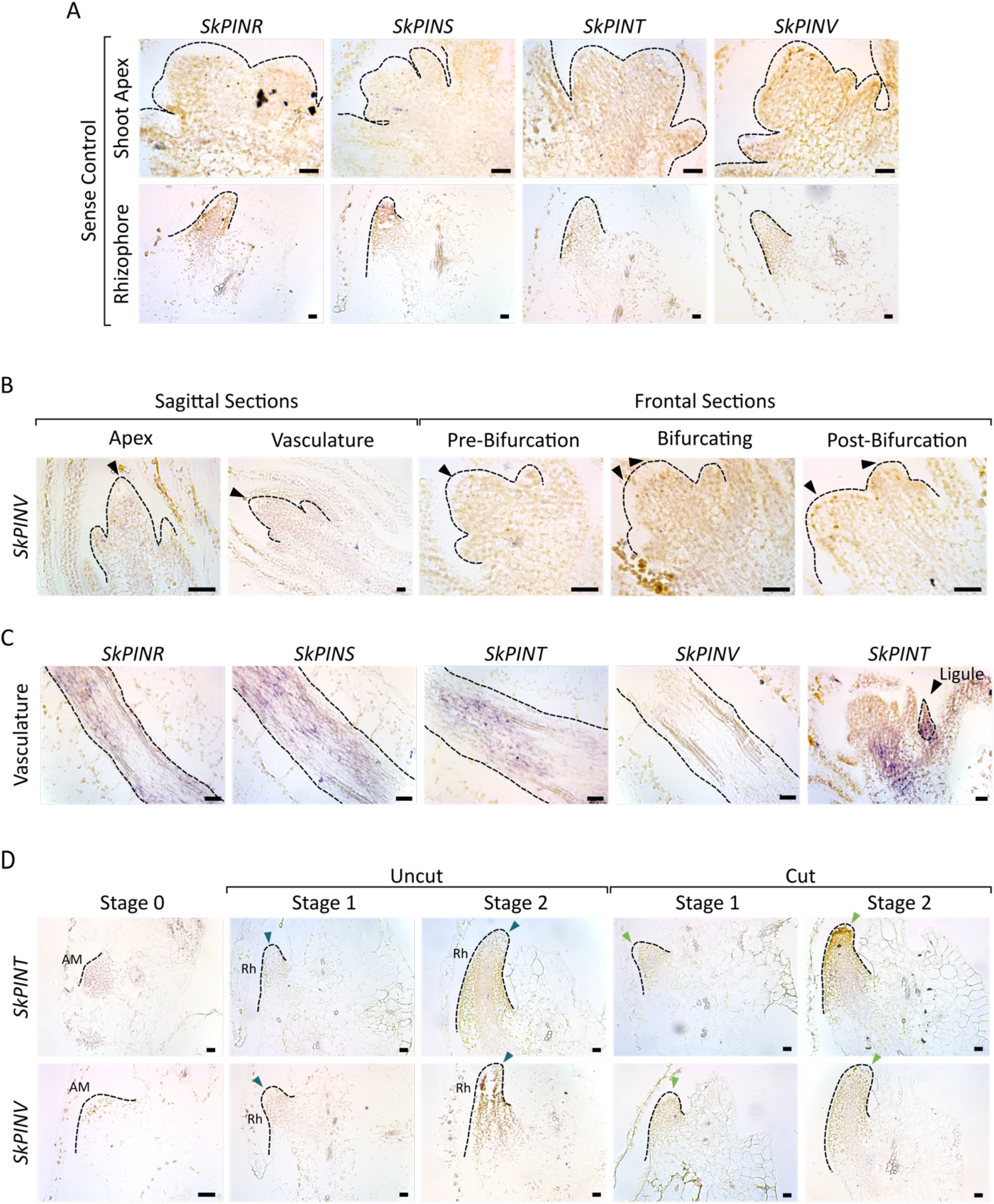
RNA *in situ* hybridisation controls and further data. (**A**) Sense controls for RNA *in situ* hybridisation of *S. kraussiana PIN* genes in the bifurcating shoot apex and rhizophore. Arrowheads = branch apex. Scale bar = 0.02 mm. (**B**) *SkPINV* expression was undetectable in frontal and sagittal sections of the shoot apex. Scale bar = 0.02 mm. (**C**) *SkPINR, SkPINS* and *SkPINT* were expressed in the vasculature. *SkPINT* was also expressed in ligules (arrowhead). Scale bar = 0.02 mm. (**D**) *SkPINT* and *SkPINV* expression in the AM and rhizophore was undetectable prior to and following apex decapitation. Blue arrowhead = rhizophore apex, green arrowhead = branch apex Scale bar = 0.02mm.

